# Matrix Stiffness Governs Fibroblast-Driven Immune Homeostasis in Gingival Tissues

**DOI:** 10.1101/2025.10.20.683155

**Authors:** Hardik Makkar, Nghi Tran, Yu-Chang Chen, Kang I Ko, Rebecca G. Wells, Kyle H. Vining

## Abstract

Periodontal disease is associated with inflamed gingival tissues and degradation of the gingival extracellular matrix (ECM), yet the role of mechanical cues is poorly understood. Gingival ECM in periodontal disease showed a loss of fibrillar collagen compared to healthy samples. We hypothesized that ECM softening in periodontal disease contributes to inflammation due to dysregulated gingival fibroblasts (GFs). A mechanically tunable hydrogel model of the gingival ECM was developed to investigate the mechano-immune crosstalk. Stiff collagen-alginate hydrogels matched the rheological properties of gingival biopsies. Human donor GFs encapsulated in these stiff hydrogels showed significantly suppressed toll-like receptor inflammatory responses compared to soft. Stiffness-dependent inflammatory responses of GFs were directed by the non-canonical NFκB pathway and epigenetic nuclear organization. The direct impact of mechanical cues on immune responses was investigated with human donor cells ex vivo by co-culture of human GFs with myeloid cells and in human gingival explants. Myeloid progenitors co-cultured with GFs in stiff hydrogels differentiated into immunomodulatory dendritic cells. Ex vivo crosslinking of human gingival tissue increased stiffness and reduced inflammatory cytokines. Gingival mechano-immune regulation provides a new avenue for biomaterials-based treatments in periodontitis.

## Introduction

Severe Periodontitis affects 19% of adults globally, causing tooth loss and increasing the risk of systemic diseases like heart disease, diabetes, respiratory infections, and adverse pregnancy outcomes^1-3^. The World Health Organization has prioritized an action plan on this condition due to its impact on health, quality of life, and healthcare costs, which exceed $40 billion USD^1^. With rising risk factors such as smoking, poor diet, and aging populations, addressing periodontitis through preventive and integrated care is crucial^3,4^. Traditional concepts of periodontal disease pathogenesis, centered around microbial dysbiosis and a resulting hyper-inflammatory host response^2,5-7^, do not adequately address how this balance is preserved in health or how it decompensates into chronic, tissue-destructive disease. The contribution of extracellular matrix (ECM) degradation and alterations in tissue mechanical properties remains understudied in gingival tissues, although it has been extensively explored in other chronic inflammatory diseases, such as rheumatoid arthritis and musculoskeletal injuries.^8^ Gingival ECM degrades in periodontitis secondary to host and microbial-derived proteases^2,4,9^. There remains a knowledge gap in how the physical microenvironment of the gingival connective tissue regulates immune homeostasis.

Gingival fibroblasts (GFs) are the resident cell type of the gingival connective tissue. GFs play a crucial role in regulating the mechano-immune axis^10-14^. They are essential both to tissue homeostasis and to the pathogenesis of periodontal disease^15-19^. GFs act as key sentinel cells in the immune response against pathogen-associated molecular patterns (PAMPs), largely through Toll-like receptor 2 (TLR2) signaling^7,10,13,20-22^. We propose that ECM breakdown and tissue softening in periodontitis triggers a switch in GF phenotype from quiescent, tissue-building cells into pro-inflammatory, matrix-degrading effectors that enhance, rather than dampen, inflammation. This establishes a destructive feedback loop in which matrix breakdown feeds a pathological immune response, which degrades more matrix and further softens the surrounding tissue^10-12,14,23^. These mechanical disturbances, in turn, have the potential to impact the nucleus (epigenome and genome) and the transcriptional control of immune responses^23^.

We investigated the mechanobiology underlying GFs’ pro-inflammatory phenotypes in periodontitis using a tunable 3D gingival ECM-mimicking hydrogel system^24-26^. We determined how matrix stiffness regulates GFs’ response to TLR2-signaling. In a stiff, healthy matrix, mechanical licensing of GFs dampens their response to TLR2-mediated inflammation. We propose that increased matrix stiffness may restore health to the gingival tissues, which we demonstrated by reduced inflammatory cytokines in human gingival explants treated with transglutaminase. Overall, these results suggest matrix stiffening may be able to restore health in periodontitis.

## Results and Discussion

### Collagen fiber organization in gingival connective tissue

Second Harmonic Generation (SHG) imaging was used to examine the three-dimensional collagen fiber structure in healthy gingival connective tissue. Whole-mount imaging of gingival connective tissue permitted a layer-by-layer examination of collagen fiber orientation in detail **(Figure 1A-D, S1A)**, represented as a heatmap. The heatmap depicts the distribution of collagen fiber pixel percentage counts at various angles (x-axis) and Z-layers (y-axis), with color intensity indicating count (red for highest counts, pink for lowest) **(Figure 1D)**. Although there is a general trend for angular orientations, the exact distribution changes dramatically with depth. In the central portion, collagen fibers are largely oriented within a restricted angular range, centered between 45-90 degrees ***(Figure 1D)***. This is suggestive of a highly organized and dominant fiber orientation^27^ within the bulk of the connective tissue, as examined previously^28^. In contrast, superficial and deeper Z-layers (towards the top and bottom of the heatmap) tend to have fibers oriented between 0-45 degrees and 135-180 degrees. The presence of distinct, high-density bands in the middle Z-layers, surrounded by regions of lower density, is indicative of a stratified arrangement of collagen fibers in the gingival tissue^27,29^. This finding indicates that collagen fibers are neither uniformly distributed nor oriented throughout the entire thickness but rather exhibit specific preferred orientations and densities at different depths, presumably reflecting the heterogeneous mechanical and structural requirements of the gingival tissue at each level^30^. Further, these findings are consistent with classic ultrastructural observations describing two principal connective tissue patterns in human gingiva.^27^

**Figure 1.**
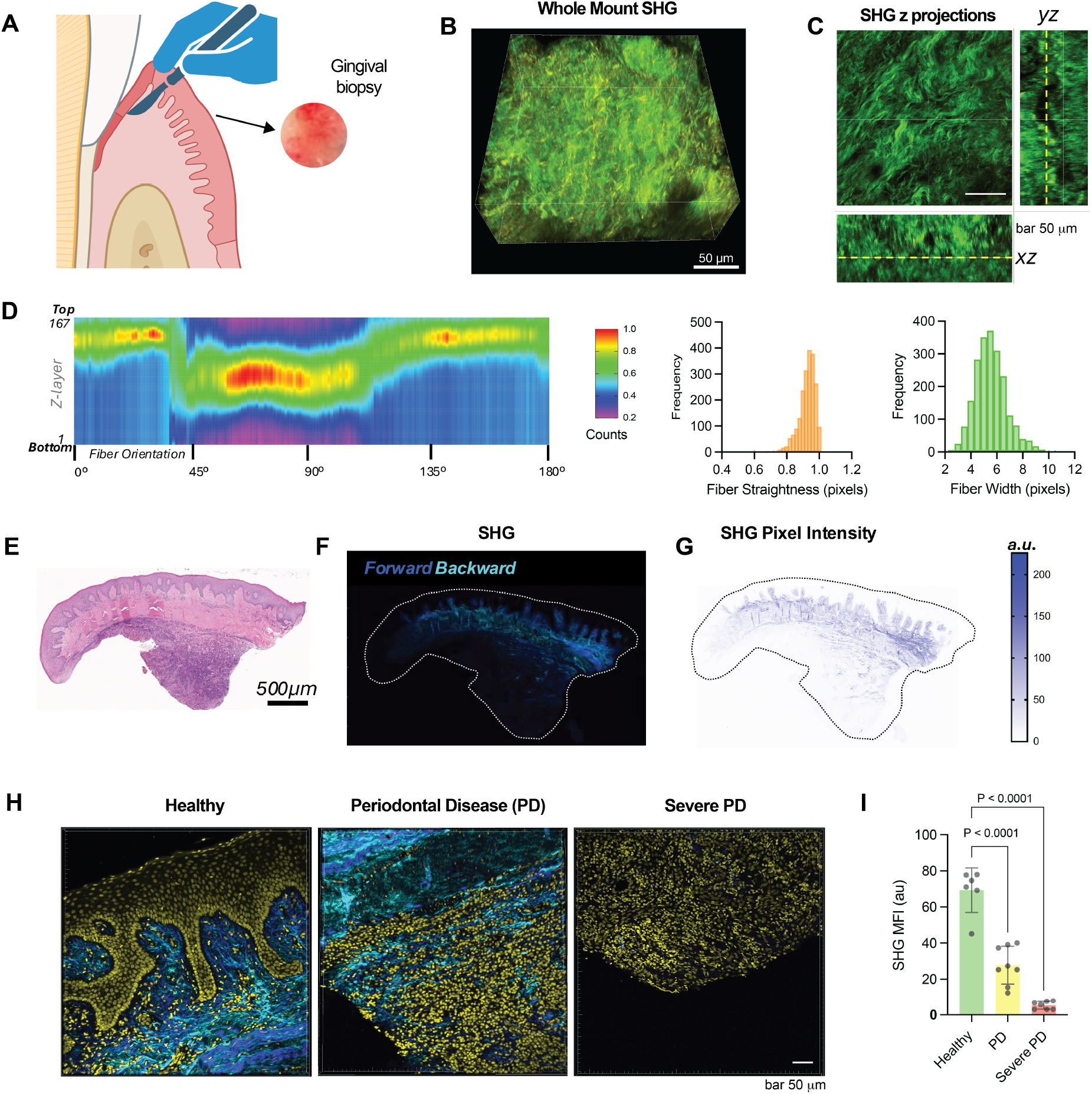
Periodontitis is characterized by a progressive loss of fibrillar collagen organization and density in the human gingival extracellular matrix (ECM). **(A)** Schematic illustrating the experimental workflow for analyzing human gingival ECM. The workflow involves the acquisition of a human gingival tissue biopsy followed by whole-mount Second Harmonic Generation (SHG) imaging for high-resolution, label-free analysis of collagen architecture. **(B)** Representative whole−mount SHG image of a healthy gingival connective tissue sample. Collagen fibers (green) are clearly visible. **(C)** SHG z−projection images (XY, YZ, and XZ planes) demonstrating the organization of collagen fibers through the tissue depth. **(D)** Heatmap showing the distribution of collagen fiber orientation as a function of Z−layer depth, corresponding histograms quantify key collagen structural parameters (straightness and width) derived from the SHG data. **(E)** Hematoxylin and Eosin (H&E) staining of a representative human gingival tissue section. **(F)** SHG forward and backward signal imaging of the tissue section outlined in (E), revealing the collagen distribution. **(G)** SHG pixel intensity map across the section in (F), providing a quantified spatial representation of collagen content. **(H)** Representative SHG micrographs of gingival connective tissue from Healthy, Periodontal Disease (PD), and Severe PD sites. **(I)** Quantification of the collagen density by measuring the Mean Fluorescence Intensity (MFI) of the SHG signal across the three disease stages. Data are shown as mean ± SD. P-values from statistical tests are indicated.

### Periodontal disease is linked with pathological changes to the extracellular matrix

Multimodal imaging, followed by quantitative analysis, was performed to understand the structural organization of collagen fibers in healthy and diseased gingival connective tissue (**Figure 1E,F)**. SHG images of healthy sites **(Figure 1F-H, S1B)** showed a robust, orderly collagen network. The images reflected dense, uniformly oriented fibrils with strong SHG signal intensity, implying the integrity and well-maintained state of collagen structures. There was little to no apparent immune cell infiltration, consistent with an inflammation-free status. Quantification of collagen fibers in healthy tissue exhibited a bimodal distribution in the orientation histogram with two major peaks at ±50−55º, showing two preferred directions and a well-organized framework (**Figure S1C)**. This finding implies two large populations of collagen fibers oriented almost orthogonally, creating an ordered, anisotropic, mechanically strong network. In sharp contrast, SHG images of pathological regions showed drastically changed tissue structure. These images showed a strong decrease in coherent SHG signal, appearing weakened and fragmented, indicating collagen degradation and disorganization (**Figure 1F-I, S1 B,C)**. Instead of well-defined fibrils, collagen was dispersed or amorphous. These images also showed heavy immune cell infiltration. Quantitative angular analysis of diseased tissue **(Figure S1 B,C)** corroborated these observations, showing a different and less organized collagen fiber distribution. While a diffuse bimodal tendency could still be seen, the peaks in diseased tissue were broader and weaker, revealing increased dispersion and randomization of orientations instead of discrete, aligned bundles. This distribution signifies a loss of preferential alignment, resulting in a disorganized and more isotropic collagen network in the diseased state, pointing to disruption of tissue integrity. A prominent trough with decreased frequency of fibers was found within ±20−25º in healthy tissue, complementing the specific directionalities and organization characteristic of gingival tissue. In diseased tissue, the trough was shallower, indicating a greater proportion of randomly oriented fibers over a wider range of angles **(Figure S1C)**. This diffuse distribution corresponds to large-scale collagen degradation and the disruptive effect of inflammatory cells seen in SHG images.

These imaging results align with prior histological and molecular studies demonstrating that periodontal inflammation drives extensive collagen degradation and remodeling of the gingival ECM. Experimental models and human biopsies consistently show a significant reduction in collagen area and altered fibril architecture during disease^29^. Periodontal pathogens and their secreted cysteine proteases not only degrade collagen directly but also induce gingival fibroblasts and epithelial cells to upregulate matrix metalloproteinases (MMP-1, MMP-2, MMP-3, MMP-9) while suppressing their inhibitors (TIMPs), amplifying host-mediated collagenolysis^31^. Immunohistochemical detection of collagenase-generated neoepitopes in chronic periodontitis further confirms active fibril cleavage by host enzymes. Collectively, the loss of fiber alignment and diminished SHG signal observed here reflect this enzymatic breakdown and imbalance between MMPs and TIMPs, leading to a weakened, disorganized ECM^32-34^. These structural and molecular alterations together highlight collagen degradation as a central mechanism driving the loss of connective tissue integrity in periodontal disease.

### Gingival ECM hydrogel models the rheological properties of human gingival tissue

SHG heatmaps generated from the gingival ECM hydrogels demonstrated their ability to replicate key aspects of collagen organization found in healthy gingival tissue **(Figure 2A-C)**. Hydrogel samples displayed a clear and significant band defined by high collagen fiber numbers, strikingly similar to the central high-density zone of native gingiva. Collagen fibers within these hydrogel bands had a strong preferential orientation, mainly concentrated between 45-135 degrees, thus mimicking the main angular directionality seen in healthy gingival tissue **(Figure 1D,2C)**. Stress relaxation tests were conducted to assess the time-dependent viscoelastic properties of the ionically cross-linked hydrogels. Prior studies have shown that stress relaxation in physical hydrogels arises from polymer chain rearrangements ^25,26^. Increasing the Ca^2+^ concentration enhanced stress relaxation **(Figure S2A,B)**, likely due to a higher density of ionic cross-links, which promotes more frequent bond rearrangements and accelerates stress dissipation^25,26,35,36^. The storage modulus (G′) and loss modulus (G′′) of the stiff hydrogel were very similar to those of healthy gingiva (∼2000 Pa) **(Figure 2D, S2C)**.

**Figure 2.**
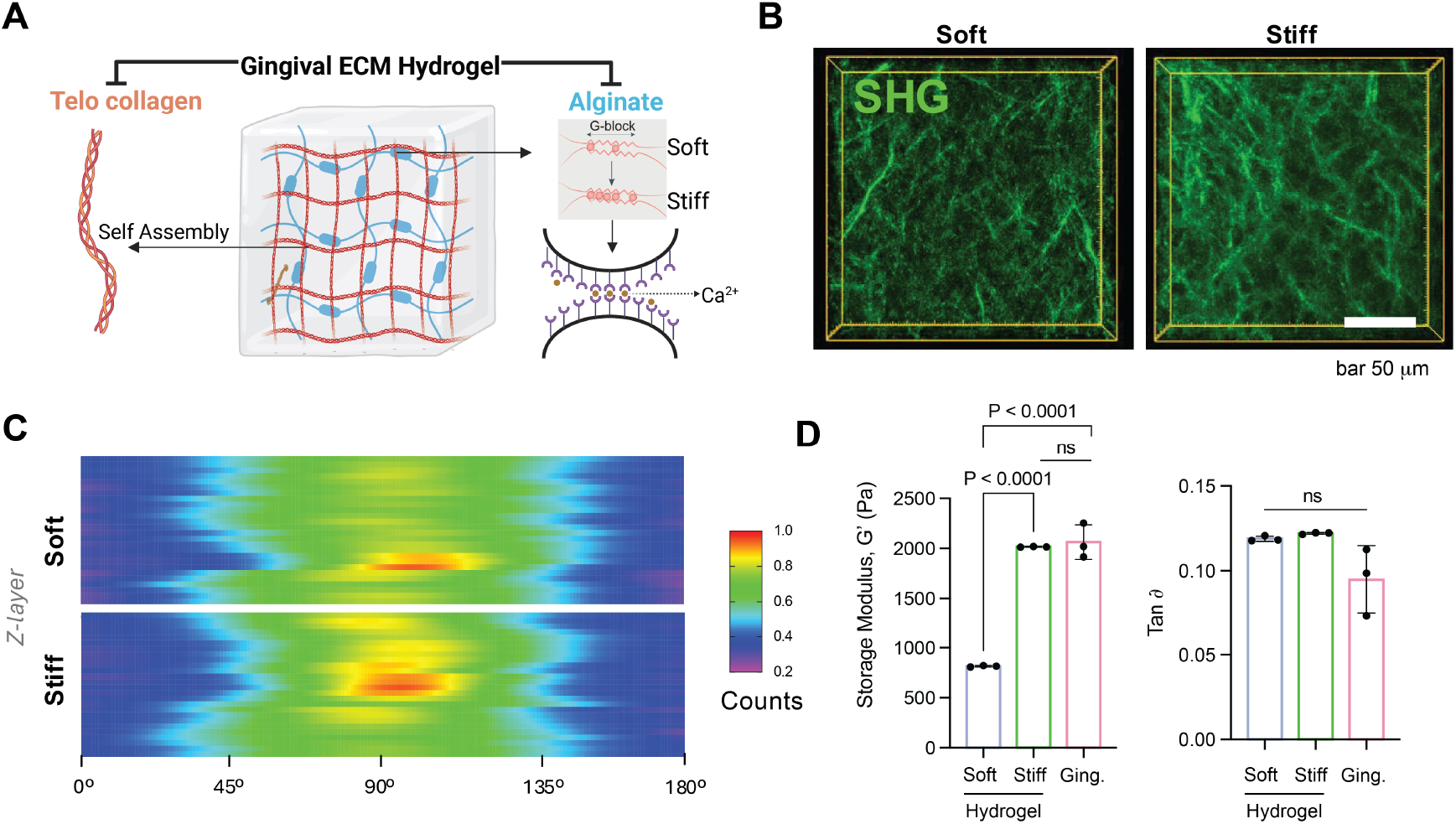
A biomimetic alginate-collagen interpenetrating hydrogel system recapitulates the tunable stiffness and viscoelasticity of human gingival ECM. **(A)** Schematic illustrating the design of the interpenetrating polymer network (IPN) hydrogel system. It is composed of self-assembled type I collagen (telo collagen) and ionically cross-linked alginate (via Ca2+ ions interacting with G-blocks). **(B)** Representative Second Harmonic Generation (SHG) images showcase the fibrillar collagen architecture within the soft and stiff hydrogel formulations. **(C)** Heatmap analysis of collagen fiber orientation distribution throughout the z−depth of the soft and stiff hydrogels. **(D)** Rheological characterization of the developed hydrogels compared to native healthy human gingival tissue. Bar graphs show the Storage Modulus (G′) and Tan Delta (Tanδ) (ratio of loss modulus G′′ to G′), representing viscoelasticity. Scale bars are as indicated. Data are shown as mean ± SD. P-values from statistical tests are indicated.

### Stiff ECM hydrogels upregulate matrisome pathways in gingival fibroblasts

Gingival fibroblasts (GFs) were isolated from healthy donor tissue and then screened for the expression of mesenchymal stem cell (MSC) surface markers using flow cytometry. GFs had a positive MSC marker expression profile. Specifically, a very high percentage of the cell population was positive for CD105, CD73, CD90, and CD44, as indicated by the clear changes in fluorescence intensity and the population clustering seen in the flow cytometry plots **(Figure S2D)**. Next, GFs were encapsulated in soft and stiff gingival ECM hydrogels. Pathway enrichment analysis of upregulated genes from bulk RNA sequencing data identified enrichment for extracellular matrix (ECM) synthesis and organization processes. Transcriptional programs related to collagen synthesis and maturation were enriched in stiff hydrogels, including the Assembly of Collagen Fibrils, Collagen biosynthesis and modifying enzymes, Collagen formation, and Collagen chain trimerization. In addition, the ECM organization was the pathway with the largest number of associated genes (Counts ≈ 200). The enrichment of the ECM proteoglycans pathway suggested a global biosynthetic response in stiff ECM hydrogels. Taken together, these findings illustrate that the mechanical stiffness of the 3D matrix promotes fibroblasts to adopt an active, matrix-synthesizing phenotype, which may help preserve tissue integrity **(Figure 3A)**. In stiff hydrogels, GFs upregulated genes reflective of a homeostatic and matrix-maintaining phenotype, typical of healthy gingival connective tissue. Notably, genes for important structural collagen constituents such as COL5A3, COL4A2, COL6A5, and COL4A1 were upregulated, pointing to active synthesis of a stable ECM **(Figure 3B, S3A)**. The upregulated expression of TIMP3, a matrix metalloproteinase inhibitor, together with TGFB2 and TGFB3 (regulators of ECM synthesis), further corroborates a transcriptional program favoring ECM deposition, maturation, and resistance to degradation. Genes such as PDK4 and CALCRL were also upregulated in GFs cultured in stiff hydrogels, demonstrating metabolic and signaling adaptations important for tissue homeostasis **(Figure S3A)**. These results were consistent with the organized collagen architecture and strong mechanical properties of native healthy gingiva^37^.

**Figure 3.**
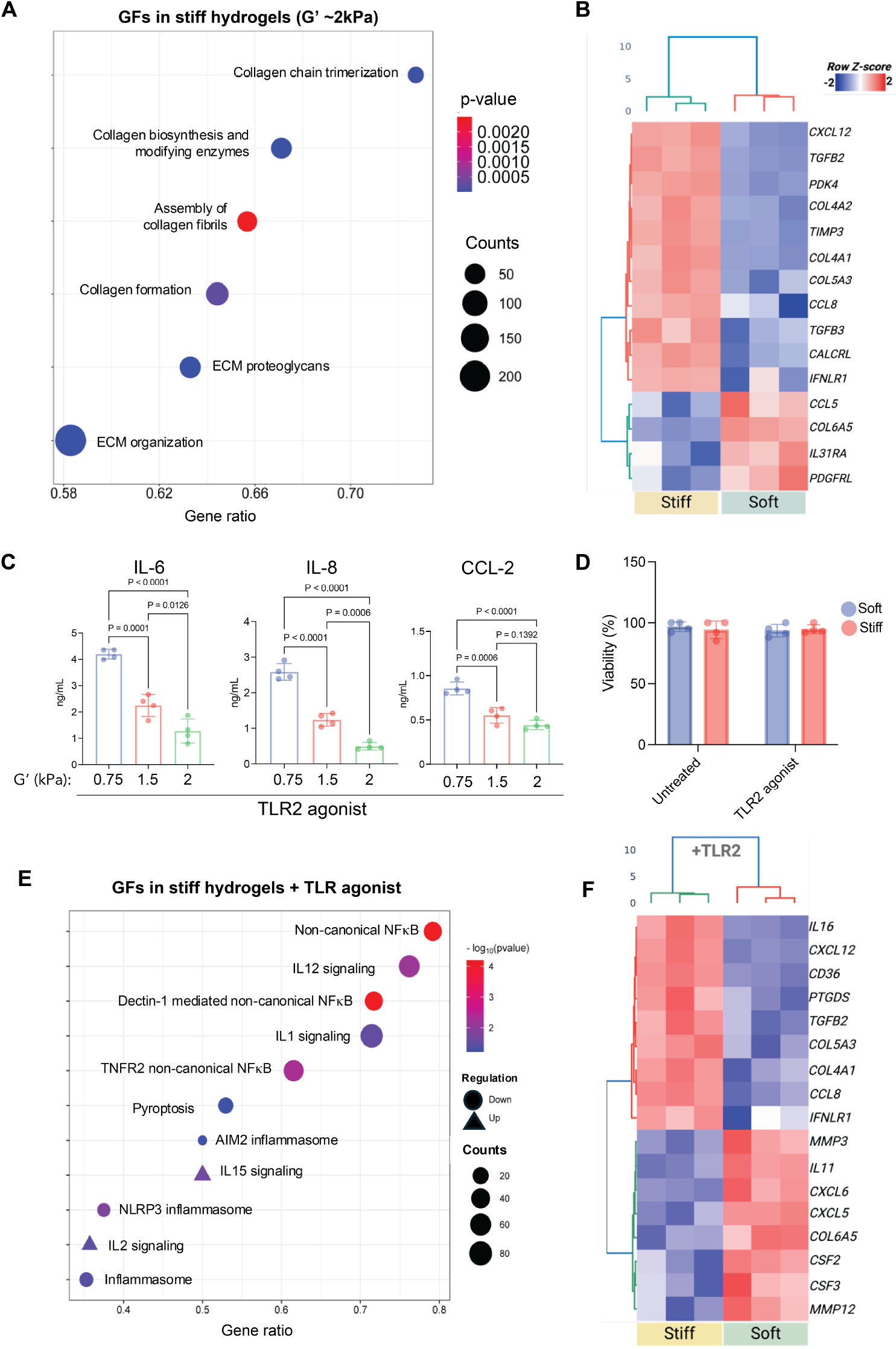
Matrix stiffness drives a pro-anabolic phenotype, while reduced stiffness supports TLR2-mediated inflammatory responses in human GFs. **(A)** Bubble plot showing the gene set enrichment of pathways significantly upregulated in GFs cultured on stiff hydrogels (G′∼2kPa) compared to soft hydrogels (G′∼0.75kPa). Pathways are predominantly related to ECM production and organization. The gene ratio (x-axis) is the proportion of differentially expressed genes in a pathway. Bubble size indicates the count (number of genes in the pathway), and the color scale represents the p-value (Benjamini-Hochberg corrected) for enrichment. **(B)** Heatmap illustrating the relative expression of differentially expressed genes in the stiff versus soft hydrogel environments. **(C)** ELISA quantification of secreted pro-inflammatory mediators IL-6, IL-8, and CCL-2 from GFs cultured in hydrogels of varying stiffness (G′: 0.75, 1.5, and 2kPa) following TLR2 activation. **(D)** Cell viability assay showing that neither hydrogel stiffness nor TLR2 agonist stimulation significantly impacted GF viability over the culture period. **(E)** Bubble plot showing the gene set enrichment of inflammatory signaling pathways regulated in GFs cultured in stiff hydrogels compared to soft, both with TLR2 agonist stimulation. Bubble size corresponds to the count, and color scale indicates the −log10(p value) of the pathway. Circles indicate downregulation, and triangles indicate upregulation in stiff hydrogels. **(F)** Heatmap showing the relative expression of differentially expressed genes in GFs cultured in soft versus stiff hydrogels, following TLR2 agonist stimulation. Data are presented as mean ± SD; P-values from statistical tests are indicated.

### TLR2 mediates Inflammatory responses in gingival ECM hydrogels

The Toll-like receptor 2 (TLR2) in gingival fibroblasts recognizes a wide spectrum of bacterial components, especially those from abundant Gram-positive and Gram-negative bacteria in periodontal biofilms^14,18,22,38^. TLR2 shows a broad recognition profile relevant for the initiation and maintenance of the chronic inflammatory condition in gingival fibroblasts^14^, whereas TLR4 recognizes Gram-negative bacterial lipopolysaccharide (LPS).^39^ Soft and stiff gingival ECM hydrogels were challenged with a TLR2 agonist to determine whether stiffness regulates fibroblast inflammatory responses. Gene expression of the TLR2 pathway was not significantly modulated by matrix stiffness, besides downregulation of CD14 **(Figure S2F)**. ELISA analysis demonstrated a dramatic, stiffness-dependent reduction of secreted pro-inflammatory cytokines. Fibroblasts within soft 3D secreted increased amounts of the cytokine IL−6 and the chemokines IL−8 (CXCL8) and CCL2 (MCP-1) in response to TLR2 stimulation **(Figure 3C, S2E)**. In contrast, increasing the stiffness of the surrounding hydrogel resulted in the progressive decline of the TLR2-mediated pro-inflammatory cytokine secretion. Fibroblasts cultured in the stiffest (2kPa) 3D matrices produced the lowest amount of IL−6, IL−8, and CCL2. These results show that mechanotransduction in a 3D tissue context is an important regulator of the fibroblast inflammatory secretome^40,41^. Higher stiffness in the surrounding ECM induced a host-protective cellular phenotype, allowing fibroblasts to limit their inflammatory response to microbial stimuli and promote tissue homeostasis.

Results from bulk RNA seq analysis indicate that GFs in soft hydrogels showed upregulation of genes commonly linked to inflammation, immune cell recruitment, and tissue breakdown, mirroring the pathologic alterations of diseased gingiva. Chemokines like CXCL12, CSF3, CCL8, CXCL6, and CXCL5 were highly upregulated in soft gels, suggesting strong chemoattractant signaling for immune cells characteristic of periodontal inflammation **(Figure 3E,F,S3B)**. Additionally, matrix metalloproteinase genes MMP3 and MMP12, disintegrin and metalloproteinase with thrombospondin motifs ADAMTS14, were upregulated in soft gels, pointing to ongoing ECM breakdown^14^. Overall, the integrated mechanotranscriptomic and inflammatory challenge analysis illustrates that the gingival ECM hydrogel system is consistent with the divergent transcriptional responses of gingival fibroblasts in healthy versus diseased gingiva. Stiff hydrogels promote a gene expression profile that actively maintains ECM integrity and regulates inflammation. Conversely, soft hydrogels promote a strong pro-inflammatory and degradative gene signature, closely approximating the disease state.

### Gingival fibroblasts exhibit morphological responses to matrix stiffness

In soft ECM hydrogels, the gingival fibroblasts displayed a characteristic, highly spread morphology with pronounced uniaxial protrusions **(Figure 4A)**. These cellular extensions were often seen to be tipped by lamellipodial blebs, reflecting active membrane dynamics underpinned by actin polymerization and cytoskeleton-mediated probing^24^. The F-actin cytoskeleton (green) was loosely organized into discrete stress fibers that extended into the processes of the cell, reflecting an active probing nature **(Figure 4A-C)**. The cell body, although spread, remained relatively elongated, and the nucleus (blue) was distended, in agreement with cells probing a compliant environment in which tension generation could be dissipated across many dynamic adhesions. This morphological phenotype is consistent with migration and remodeling ^35^.

**Figure 4.**
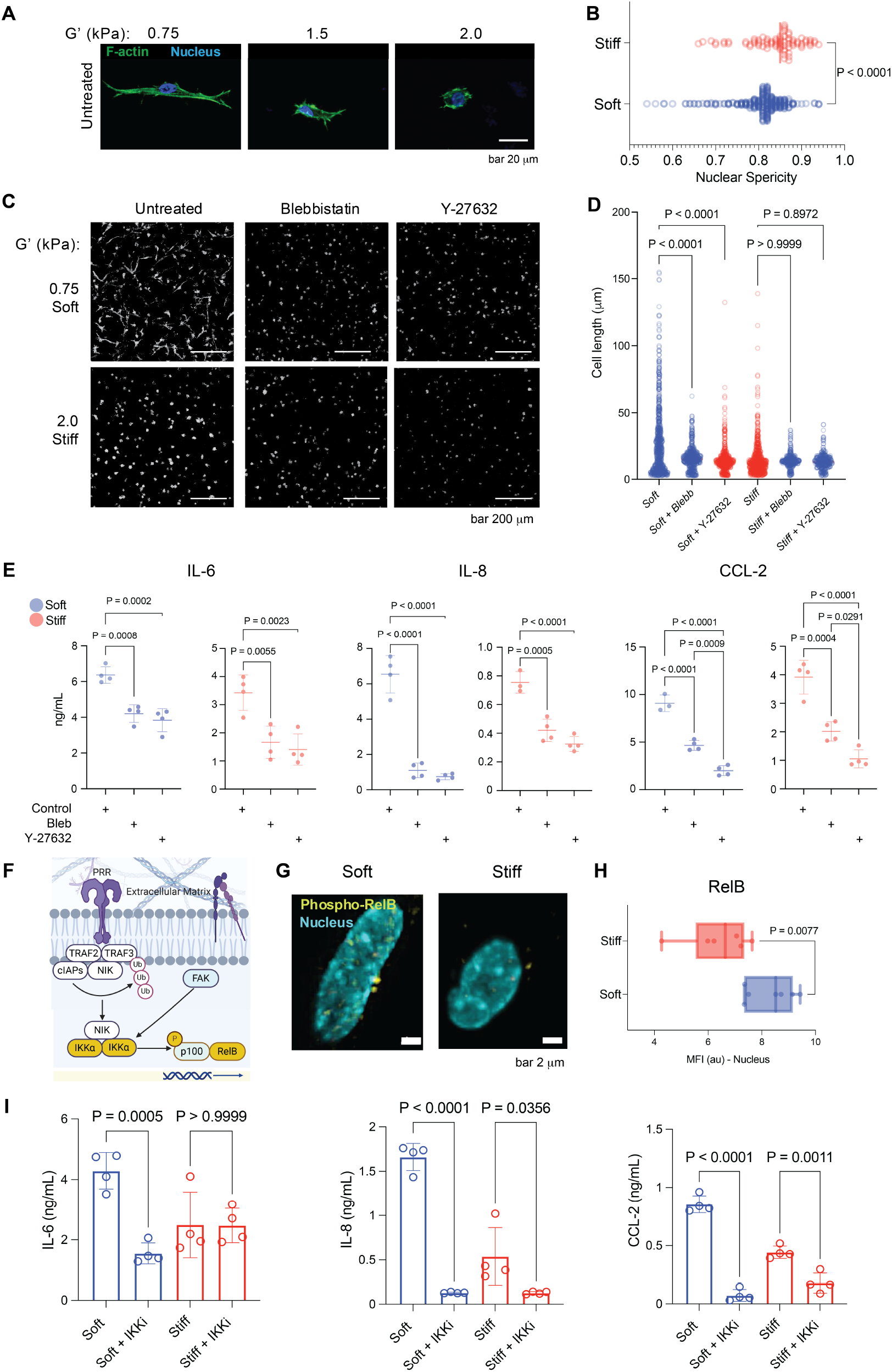
Cytoskeletal tension and the non-canonical NF−κB pathway regulate matrix stiffness-dependent inflammatory responses in gingival fibroblasts (GFs). **(A)** Representative confocal immunofluorescence images showing F-actin organization (Phalloidin, green) and nuclear morphology (DAPI, blue) in GFs cultured on hydrogels with increasing stiffness (G′: 0.75, 1.5, and 2.0kPa). **(B)** Quantification of nuclear sphericity confirms the stiffness-dependent nuclear condensation observed in GFs. **(C)** Representative images and **(D)** quantification of cell length in GFs following treatment with the non-muscle myosin II inhibitor Blebbistatin (Blebb) or the ROCK inhibitor Y-27632. **(E)** ELISA quantification of IL−6, IL−8, and CCL−2 secretion from TLR2-activated GFs treated with cytoskeletal inhibitors. **(F)** Schematic representation of the proposed mechanism: ECM stiffness and TLR2 signaling converge to regulate non-canonical NF−κB signaling in GFs. **(G)** Representative immunofluorescence images and (H) quantification of the Mean Fluorescence Intensity (MFI) of nuclear-localized phosphorylated RelB (Ser552) in cells cultured in soft versus stiff hydrogels. **(I)** ELISA quantification of IL−6, IL−8, and CCL−2 secretion from TLR2-activated GFs cultured in soft or stiff matrices with or without IKK inhibition. Scale bars are as indicated. Data are shown as mean ± SD. P-values from statistical tests are indicated.

By contrast, the spreading of gingival fibroblasts in stiff hydrogels was significantly hindered. The cells displayed a more compact, round, or stellate morphology. The formation of uniaxial extensions and lamellipodial blebs was greatly reduced. While F-actin (green) was still visible, it was organized into a more compact and highly tensed structure within the limited cellular space **(Figure 4A)**. The nucleus also showed a more spherical morphology compared to cells in softer gels, reflecting the altered cytoskeletal tension and overall cellular confinement without any impact on cellular viability **(Figure 4B, 3D)**. Active spreading and probing protrusions on soft hydrogels are indicative of a phenotype usually seen with migration and exploration of new or compliant tissue^42^. In stiff hydrogels, however, the constrained spreading and more compact morphology are indicative of a cell-matrix signaling ^43^.

### Actomyosin Contractility Determines Fibroblast Morphology and Inflammatory Response in a Stiffness-Dependent Manner

The intrinsic mechanosensitivity of GFs was also verified by manipulating their actomyosin contractility. GFs encapsulated in the hydrogels were exposed to blebbistatin (a myosin II inhibitor) or Y27632 (an inhibitor of Rho-associated protein kinase (ROCK)), both key disruptors of actin cytoskeleton-generated forces^24^. Confocal micrographs in **Figure 4C** show that the clear morphological responses of GFs to matrix stiffness were greatly changed by these inhibitors. In soft ECM hydrogels, untreated GFs displayed elongated lengths, representative of their active spreading and exploratory morphology. With inhibition of actomyosin contractility, a pronounced and significant decrease in cell length was observed, with the population of fibroblasts taking on a shorter, more rounded morphology **(Figure 4D)**. This illustrates that the typical elongated phenotype of GFs in a soft environment is dependent, to a critical extent, on their inherent actomyosin contractile forces^12,44,45^. Likewise, in stiff hydrogels, although GFs were already more compact, inhibition of contractility also caused a reduction in cell length, confirming the general dependence of GF morphology on actomyosin activity across a range of matrix stiffness. To further investigate the intersection of actomyosin contractility with inflammation, GFs in both soft and stiff hydrogels were challenged with a TLR2 agonist to promote an inflammatory response. Following this, the impact of actomyosin contractility inhibition on inflammatory markers was evaluated. Strikingly, treatment with blebbistatin and Y27632 dampened the inflammatory response within the gingival fibroblasts. Most importantly, immune responses were stiffness-independent because the decrease in inflammatory markers was seen to a similar extent in both the soft and stiff hydrogels **(Figure 4E)**. This indicates that inhibition of actomyosin contractility blocked the ability of GFs to sense ECM stiffness ^24,39,46^, demonstrating that intracellular force generation is essential for mechanosensitive inflammatory signaling. Under these conditions, TLR2-driven responses are uniformly attenuated, indicating that actomyosin-mediated mechanotransduction underlies the differential inflammatory responses^45,47^.

### Mechanotransduction-induced inflammatory response in GFs is controlled by non-canonical NF-κB signaling

Other innate immune pathways, in addition to TLR2, were assessed by stimulation of GFs with agonists for Toll-like receptors 3 (TLR3) and 4 (TLR4). The data revealed a similar pattern across these TLRs. GFs in soft hydrogels responded to TLR3 and TLR4 agonists with a strong inflammatory response, with high expression of IL-6 and IL-8 **(Figure S2H)**. GFs in stiff hydrogels, however, displayed a significantly subdued inflammatory response. These data verify that the dampened inflammatory response by a stiff matrix is not an isolated event at one pattern recognition receptor but rather a more general regulatory mechanism controlling the innate immune response of GFs. For simplicity, the remaining studies focused on the mechanoregulation of TLR2-mediated inflammation.

Pathway enrichment analysis was conducted on differentially expressed genes from bulk RNA-sequencing of GFs cultured on stiff versus soft gels activated with a TLR2 agonist. The analysis, as represented by the bubble plot **(Figure 3E)**, reveals differentially regulated signaling pathways with the highest fold enrichment, giving important insight into the potential protective mechanisms of a healthy extracellular matrix. The most striking result was the strong downregulation of non-canonical NF-κB in stiff hydrogels **(Figure 3E, 4F)**. The analysis also indicated downregulation of multiple inflammasome-related pathways, which further supports a role for healthy matrix mechanics in dampening potentially harmful inflammatory cascades. Inflammasome-related pathways were upregulated, albeit with generally lower statistical significance than the highly downregulated pathways. These included IL15 signaling and IL2 signaling. The engagement of these cytokine-signaling pathways, however modest, may be a compensatory or regulatory response in the healthy-mimicking matrix to modulate the overall immune modulation^48^. To further confirm the observed downregulation of non-canonical NF-κB signaling, we examined protein levels of RelB, an important transcription factor of this pathway^49^. Immunofluorescence staining and subsequent quantification of lower RelB mean fluorescence intensity (MFI) in gingival fibroblasts encapsulated in stiff hydrogels gave direct confirmation of these molecular observations **(Figure 4G,H)**.

IKK, a kinase upstream of RelB, was inhibited following TLR2 activation of GFs in hydrogels. The secretion of IL-6, IL-8, and CCL-2 in soft hydrogels was greatly diminished, reaching levels comparable to or even below those of stiff gels. This drastic decrease shows that the activated inflammatory response in soft matrices is reliant on the activity of the IKK complex^49^. In contrast, TLR2-stimulated GFs in stiff hydrogels had dramatically lower baseline levels of these inflammatory markers, which align with our observations of dampened inflammatory signaling in a healthy mechanical environment. Critically, the IKK inhibitor had little to no further effect on the expression of IL-6 in stiff hydrogels **(Figure 4I)**. This absence of significant change further validates our hypothesis that the stiff matrix inherently downregulates the non-canonical NF-κB pathway, making it less active and hence less responsive to further pharmacological inhibition. Collectively, these functional experiments give strong evidence that the mechanically-regulated non-canonical NF-κB^49^ signaling pathway is a fundamental determinant of the gingival fibroblast’s inflammatory phenotype to mechanical cues and inflammatory stimuli in a 3D environment.

### GFs exhibit stiffness-dependent decrease in overall nuclear volume and transcriptional activation of pathways linked with the ECM-cytoskeletal-nuclear axis

Next, we examined the impact of matrix stiffness on nuclear shape. Confocal microscopy revealed that there were significant differences in the nuclear morphology of GFs embedded in soft compared to stiff hydrogels. GFs in soft hydrogels had more elongated nuclei, while GFs in stiff hydrogels had rounder nuclei. Further, quantification verified increased nuclear sphericity in GFs in stiff matrices **(Figure 4B, S4B, S4C)**. Stiffer matrices tend to induce increased cytoskeletal tension that is transmitted to the nucleus, compressing it into a more compact and spherical shape^50^. Conversely, the decreased tension on softer matrices allows for a more elongated or irregular nuclear morphology^51^. Since the nucleus is a key mechanosensitive organelle that integrates mechanical cues impinging on chromatin structure and gene expression^50-53^, these differential changes in nuclear shape point to a basic mechanism by which matrix stiffness directly modulates the phenotype of gingival fibroblasts, therefore playing a critical role in distinguishing between healthy and pathological states. To further understand this process, we conducted a targeted pathway enrichment analysis to identify major molecular cascades implicated in the mechanotransduction from ECM to the nucleus of GFs cultured in TLR2-activated stiff gingival ECM hydrogels. The bubble plot displays the most highly upregulated pathways in stiff hydrogels related to the ECM-cytoskeleton-nuclear axis **(Figure S4A)**. Analysis revealed the concerted activation of pathways outlining the intricate communication network that couples external mechanical cues to nuclear response. Integrin cell surface interactions and non-integrin membrane-ECM interactions pathways were prominently upregulated, highlighting the crucial roles played by both direct and indirect cell-ECM adhesion in sensing matrix stiffness. These interactions represent the major conduits for the relay of forces from the ECM into the cell body. Additionally, lamin interactions were upregulated, highlighting the physical linkage between the cytoskeleton and the nuclear lamina, which is critical for the transmission of physical forces to the nucleus, influencing nuclear shape and transcriptional activity^54,55^. Immunostaining for Lamin A/C showed that GFs in soft hydrogels displayed a more peripheral Lamin A/C distribution, concentrated mainly at the nuclear rim **(Figure S4B)**. This peripheral localization is typically associated with a mechanically rigid nuclear lamina, reduced nuclear deformability, and constrained chromatin organization^50,51,55,56^. In contrast, in stiff hydrogels, Lamin A/C was more diffusely distributed, extending into the nucleoplasm rather than being restricted to the periphery. Such diffuse localization reflects a more dynamic and deformable nucleus, facilitating chromatin remodeling and enhanced transmission of mechanical signals from the ECM into the nuclear interior^50,51^.

Next, we examined the role of actomyosin contractility on nuclear morphology. GFs in stiff gingival hydrogels consistently had a much smaller nuclear volume than GFs in soft gels, regardless of the presence of inhibitors of actomyosin contractility **(Figure S4C)**. This robust matrix stiffness-dependent difference shows that the mechanical properties of the extracellular milieu set a fundamental baseline or set point for nuclear volume. Within each condition of stiffness, actomyosin contractility inhibition decreased nuclear volume compared to their respective untreated controls. Notably, this decrease did not negate the intrinsic nuclear volume difference imposed by the matrix stiffness; nuclear volume in inhibitor-treated stiff gels was still much lower than in inhibitor-treated soft gels. These results demonstrate a biphasic regulatory mechanism for nuclear morphology. Extracellular matrix stiffness largely determines the baseline nuclear volume, likely through external mechanical confinement and/or stiffness-mediated influences on nuclear chromatin density^56^. Active cellular actomyosin contractility acts as a dynamic modulator of nuclear volume and shape, exerting tensile forces that work to stretch the nucleus and influence its compaction^45,51^. Inhibition of these internal forces results in a more compact, intrinsically spherical nuclear morphology. This complex interplay highlights that the nucleus is regulated by both the constant physical constraints of the ECM and the dynamic internal forces generated by the cell, which together establish its mechanosensitive state and, thereby, its functional and epigenetic landscape^53^ in periodontal health and disease.

### Inhibition of DNA methyltransferase eliminates the effect of matrix stiffness on nuclear morphology, spreading, and inflammatory response of gingival fibroblasts

Given that matrix stiffness has a profound impact on nuclear morphology and, as our pathway analysis suggests, induces epigenetic changes, we hypothesized that this would promote differential levels of nuclear condensation to maintain distinct cellular phenotypes, potentially through DNA methylation pathways^23,57-63^ **(Figure S4D)**. To test this hypothesis, we exposed gingival fibroblasts (GFs) embedded in soft and stiff hydrogels to DNA methyltransferase inhibitors (DNMTi), known for their ability to reduce DNA methylation and possibly induce chromatin decondensation. We observed a dose-dependent impact of DNMT inhibition on gingival fibroblasts embedded in stiff hydrogels. Fibroblasts in stiff gels generally exhibited higher DNMT1 nuclear expression **(Figure 5A)**, a more rounded nuclear shape, and may represent a transcriptionally repressed state. DNMT inhibition showed a pronounced increase in cellular spreading in stiff hydrogels, consistent with our hypothesis that DNA methylation pathways regulate stiffness-dependent responses in GFs **(Figure 5B, C)**. DNMT inhibition also caused a significant increase in nuclear volume for fibroblasts cultured in stiff hydrogels **(Figure 5D)**. This finding provides direct functional evidence that the more compact or condensed nuclear morphology observed in fibroblasts in stiff ECM is partly maintained by DNA methylation. An increase in nuclear volume often correlates with decondensation of chromatin, a process that may increase DNA accessibility and potentially alter gene expression patterns. This observation is consistent with current research showing that increased global DNA methylation is observed in stiffer matrices and that epigenetic processes integrate microenvironmental inputs to regulate fibroblast activation^8,64-66^. These findings strongly support our hypothesis that matrix stiffness causes nuclear changes, in part through epigenetic mechanisms of DNA methylation^58,59,62^, which affects cellular morphology and nuclear volume. Our finding that DNMT inhibition can reverse components of the GF “stiff-matrix phenotype” highlights the important role of tissue mechanics in controlling cellular epigenetic landscapes^63^ and ultimately determining cell behavior and phenotype in gingival health and disease.

**Figure 5.**
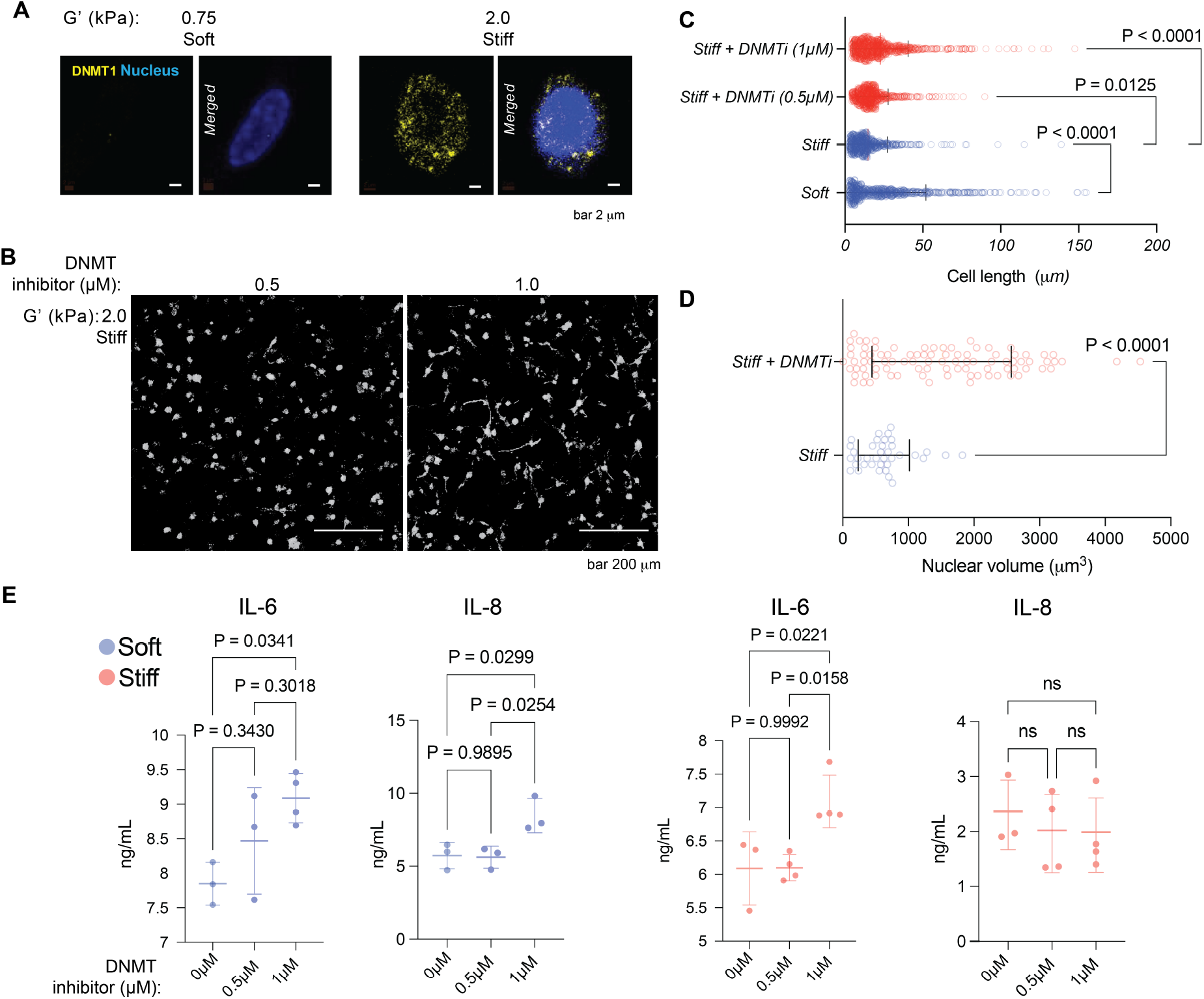
DNMT1 inhibition rescues the mechanical phenotype and restores pro-inflammatory cytokine secretion in fibroblasts on stiff matrices. **(A)** Representative confocal immunofluorescence images showing the localization of DNMT1 (yellow) and the nucleus (DAPI, blue) in gingival fibroblasts (GFs) cultured on soft (G′: 0.75kPa) versus stiff (G′: 2.0kPa) ECM hydrogels. **(B)** Representative Z-projection images of phalloidin-stained GFs encapsulated in stiff hydrogels (G′: 2.0kPa) treated with increasing concentrations of the DNMT inhibitor (0.5μM and 1.0μM). **(C)** Quantification of cell length. **(D)** Nuclear volume analysis. **(E)** ELISA quantification of IL−6 and IL−8 secretion from GFs cultured on soft or stiff hydrogels with or without DNMT inhibitor treatment. Scale bars are as indicated. Data are shown as mean ± SD. P-values from statistical tests are indicated.

We further hypothesized that this epigenetic control could directly impact the stiffness-dependent inflammatory response. To examine this, gingival fibroblasts (GFs) in stiff and soft hydrogels were treated with a DNMT inhibitor, then stimulated with a TLR2 agonist, and the secretion of the pro-inflammatory cytokines IL-6 and IL-8 was measured by ELISA. We observed that the immunomodulatory effect of matrix stiffness was completely abrogated. The stiff hydrogels, which had suppressed IL-6 secretion, now secreted significantly more IL-6, achieving concentrations indistinguishable from those secreted by GFs in soft hydrogels **(Figure 5E)**. This result is compelling functional evidence that DNA methylation plays a central role in mediating the mechanosensitive inflammatory response of gingival fibroblasts. By inhibiting DNA methyltransferases, we effectively epigenetically soften the cellular response to the stiff matrix, resulting in a pro-inflammatory phenotype. even though the bulk hydrogel stiffness was unchanged upon treatment. This is evident that mechanically induced DNA methylation is an important epigenetic mechanism^52,63-66^ by which healthy gingival ECM actively suppresses pro-inflammatory pathways. When this epigenetic control is removed, the fibroblasts lose their intrinsic anti-inflammatory ability, recapitulating aspects of the deregulated inflammatory response of periodontitis.

### Human co-culture model and gingival explant model support the mechano-modulation of inflammation in periodontitis

We next investigated whether mechanical crosslinking of the gingival extracellular matrix (ECM) may improve management of periodontitis through modulating the local immune milieu **(Figure 6A)**. We propose that an ECM crosslinker, i.e., transglutaminase, may be used to strengthen the gingival ECM. This mechanically reinforced ECM would, in turn, modulate the pro-inflammatory behaviors of resident cells (earlier in soft and degraded matrix) and promote immunological homeostasis, breaking the cycle of chronic inflammation and tissue breakdown typical of periodontal disease. Collagen has nonhelical ends (telopeptide regions), which are even more pronounced in degraded matrix, with accessible glutamine and lysine residues^25^. Transglutaminase acts on these sites, covalently linking the glutamine side chains to lysine via an acyl transfer reaction^67^. This causes crosslinking of adjacent collagen fibrils or collagen with adjacent ECM proteins like fibronectin and laminin^68-70^.

**Figure 6.**
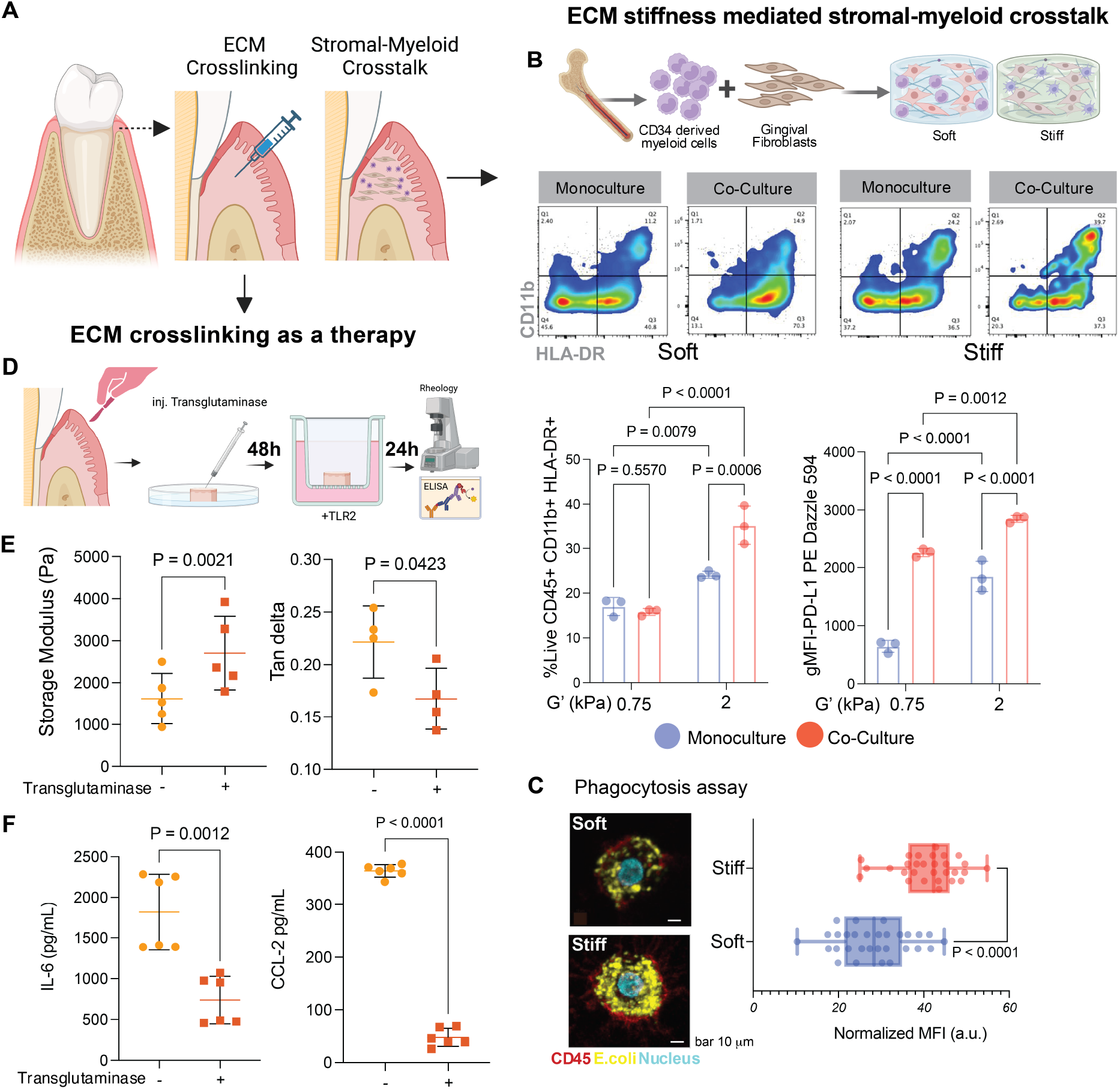
Mechano-modulation of Periodontal Inflammation via ECM stiffness. **(A)** Schematic illustrating two human-relevant models developed to investigate and therapeutically target ECM stiffness in periodontal inflammation. The left panel shows the ex vivo human gingival explant model, where ECM stiffness is actively increased using enzymatic collagen crosslinking (Transglutaminase), mimicking a therapeutic intervention to treat tissue softening in periodontal disease. The right panel illustrates the immune-competent in vitro co-culture model, utilizing mechanically tunable ECM hydrogels to study stiffness-dependent stromal–myeloid crosstalk between GFs and CD34+-derived myeloid cells. **(B)** In vitro ECM stiffness-mediated stromal–myeloid crosstalk. Schematic depicts the 3D co-culture of human CD34+ hematopoietic stem cell (HSC)-derived myeloid precursors and GFs within soft (G′: 0.75kPa) or stiff (G′: 2.0kPa) hydrogels. Representative flow cytometry plots and quantification (bottom panels) of myeloid cells (live, CD45+ gated), percentage of CD11b+HLA−DR+ cells (left), and PD−L1 expression (right, gMFI). **(C)** Representative micrographs of soft and stiff ECM-derived dendritic cells after incubation with pHrodo™ *E. coli* bioparticles, with quantification of phagocytosis by normalized mean fluorescence intensity (MFI) of particle uptake. **(D)** Experimental workflow detailing the use of the ex vivo human gingival explant model for translational validation, involving Transglutaminase injection, subsequent TLR2 stimulation, and rheological/ELISA analysis. **(E)** Rheological characterization confirms that Transglutaminase treatment successfully increases the ECM stiffness (Storage Modulus, G′) and decreases Tanδ of the human gingival tissue. **(F)** ELISA quantification of IL−6 and CCL2 secretion from TLR2-stimulated explants. Scale bars are as indicated. Data are shown as mean ± SD. P-values from statistical tests are indicated.

We developed a clinically relevant model to examine the relationship between ECM mechanics and immune cell function. First, a mechanically tunable gingival ECM hydrogel mimicked stromal-immune cell interaction by co-culture of human GFs and CD34+ hematopoietic stem cell-derived myeloid progenitors **(Figure 6B)**. This strategy was pursued to circumvent the confounding effects of previous antigen exposure intrinsic to peripheral blood monocytes, so that de novo differentiation could be more clearly evaluated^60,71,72^. Naive myeloid cells were then encapsulated in soft or stiff ECM hydrogels, either as monoculture or in co-culture with GFs. We then examined the differentiation, functional potential, and immunomodulatory phenotype of the stem cell-derived myeloid cells to discern the integrated impact of fibroblast-derived cues and mechanical signals. The combination of a stiff matrix and co-culture with GF promoted differentiation of the monocytes to functionally mature DCs with a higher population of HLA−DR+ CD11b+ cells (∼40%) **(Figure 6B)**. To determine if these phenotypic changes were coupled to enhanced immunological function, we measured the phagocytic activity of the cells against pathogen mimetics. Immune cells grown in stiff hydrogels had a significantly greater phagocytic activity than those cultured in more compliant gels, as measured by the internalization of pH-sensitive fluorescent E. coli particles **(Figure 6C)**. This indicates that a mechanically stiff environment, which is a hallmark of healthy gingival tissue, supports the development of functionally competent antigen-presenting cells^47,73,74^. We explored whether these DCs were also armed with immunomodulatory function by analyzing the expression of the immune checkpoint ligand PD-L1. Both matrix stiffness and GF co-culture were potent inducers of PD-L1. In immune cell monoculture, stiffness alone resulted in a notable increase in PD-L1 expression. Yet, co-culture with GFs induced a further upregulation in both soft and stiff matrices **(Figure 6B, S4F)**. Highest PD-L1 expression was seen on cells from the stiff co-culture condition, suggesting a robust synergy between mechanical and paracrine signaling in dictating the cells’ regulatory potential. Overall, our data show that GFs in a mechanically stiff microenvironment orchestrate a multifaceted differentiation program. Not only do they push myeloid precursors into becoming functionally competent, phagocytic DCs, but they also concurrently equip them with a strong immunomodulatory phenotype in the form of high PD-L1 expression, suggesting a mechanism for the maintenance of immune homeostasis in healthy tissue^75,76^.

Next, we tested our hypothesis in an ex vivo human gingival explant model, which maintains the native tissue structure, cellular heterogeneity, and ECM composition **(Figure 6D)**. This human tissue model avoids inter-species differences in immunology and matrix biology of animal models, which consequently offers more clinically relevant data^15,17^. We employed this model to evaluate the therapeutic potential of our crosslinking approach. Delivery of transglutaminase (Tg) to the explants successfully remodeled the mechanical properties of the tissue. Rheological measurements showed that Tg-treated tissue had a greater storage modulus (G′) and a lower tan delta (Tan δ=G′′/G′) value **(Figure 6E)**. Significantly, this mechanical transformation/change was accompanied by a reduced inflammatory response. When challenged with a TLR2 agonist to mimic a bacterial challenge, the Tg-treated explants displayed a significant reduction in the release of key inflammatory mediators **(Figure 6F, S5)**. Secretion of the pro-inflammatory cytokine IL-6 and monocyte chemoattractant CCL2 was reduced in the stiffened tissue when compared to untreated controls. This result demonstrates that by restoring the mechanical integrity of the extracellular matrix (ECM), it is possible to effectively uncouple the pathogenic stimulus (TLR2 activation) from its downstream inflammatory cascade. The lowering of CCL2 is especially relevant, as it would restrict the recruitment of further inflammatory monocytes, thereby disrupting a critical positive feedback loop that perpetuates chronic inflammation in periodontitis^77,78^.

Overall, findings from these complementary humanized platforms offer strong evidence that the mechanical properties of the ECM are a key regulator of immune function in the gingiva. The use of such systems is not only valuable to basic mechanobiology research but also as effective preclinical platforms for biomaterials development and validation. Our data suggest that ECM crosslinking as a potential therapeutic approach to periodontitis is worthy of further investigation, as we show that restoring tissue stiffness is sufficient to flip the local immune environment from pro-inflammatory to homeostatic and pro-resolution.

## Conclusion

This study provides compelling evidence that the mechanical microenvironment of the gingival connective tissue is an active and critical regulator of gingival fibroblast (GF) phenotype and immune homeostasis. Through the development and application of a tunable 3D gingival extracellular matrix (ECM)-mimicking hydrogel system, we have identified a TLR2-mediated mechano-immunological checkpoint, demonstrating how matrix stiffness programs GFs to orchestrate either a homeostatic or a pathological response. Our findings reveal that stiff matrices promote a GF phenotype characterized by enhanced ECM synthesis, organized nuclear morphology, and a robust suppression of pro-inflammatory responses, largely mediated by the non-canonical NF-κB pathway and influenced by DNA methylation. Conversely, soft, disease-mimicking hydrogels drive GFs towards a pro-inflammatory, matrix-degrading phenotype, recapitulating key transcriptional signatures observed in human periodontitis fibroblasts. Furthermore, we have elucidated the intricate crosstalk between GFs and myeloid cells, showing that GFs in a mechanically stiff microenvironment synergistically promote the differentiation of functionally competent, phagocytic dendritic cells (DCs) while simultaneously equipping them with a strong immunomodulatory phenotype via PD-L1 expression. This multifaceted regulatory program underscores a novel mechanism for the maintenance of immune homeostasis in healthy gingival tissue. In summary, our work establishes a paradigm shift in understanding periodontal disease pathogenesis, moving beyond traditional microbial and hyper-inflammatory host response models to highlight the central role of the physical microenvironment ***(Figure 7)***. This research not only confirms the validity of our experimental system but also emphasizes matrix stiffness as an essential, context-dependent modulator of fibroblast phenotype in periodontal health and disease, opening new avenues for therapeutic intervention. The insights gained from this study lay a robust foundation for several exciting avenues of future research, elucidating the precise mechanistic link between DNA methylation and NF-κB signaling, investigating the role of nuclear mechanics and chromatin remodeling in GF mechanotransduction, and exploring the translational potential of mechano-immune therapies through biomaterial-based strategies to restore healthy tissue stiffness. Future studies will focus on exploring the heterogeneity of gingival fibroblasts using our 3D hydrogel model to identify novel therapeutic targets within specific cellular subsets and conducting *in vivo* validation and preclinical models to translate these findings for the treatment of periodontitis.

**Figure 7.**
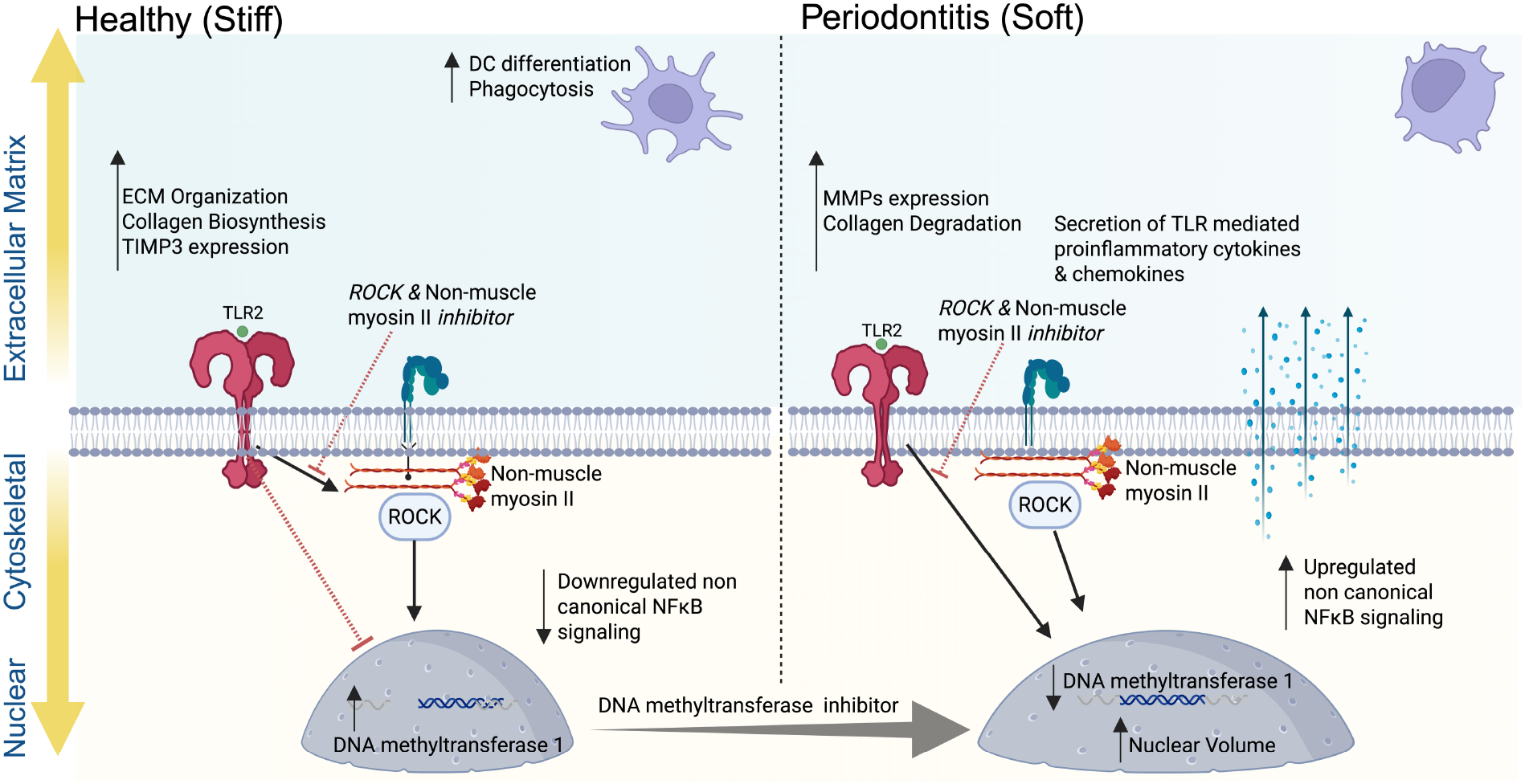
The ECM-Cytoskeletal-Nuclear Axis Regulates Periodontal Inflammation and Fibroblast Phenotype via DNMT1 and Non-Canonical NF−κB Signaling.

## Acknowledgements

This work was supported by the National Institute of Dental and Craniofacial Research (NIDCR) through a training grant to the Center for Innovation & Precision Dentistry (CiPD) (R90DE031532 to Hardik Makkar). Additional support was provided in part by the Collaborative Research Grant from the Institute for Regenerative Medicine in the Perelman School of Medicine and the School of Dental Medicine at the University of Pennsylvania (Vining). This study was also partially supported by the National Institute of General Medical Sciences (NIGMS) (R35GM157079, Vining and Chen) and by a Graduate Research Fellowship from the National Science Foundation (No. DGE-2236662, Nghi Tran). We thank Gordon Ruthel at the Penn Vet Imaging Core for confocal microscopy and SHG imaging assistance. Confocal microscopy was performed on an instrument purchased with support from an NIH Shared Instrumentation Grant (S10 OD032305-01A1). We also thank Jennifer Murray at the Children’s Hospital of Philadelphia Flow Cytometry Core for assistance with flow cytometry.

## Author Contributions

**CRediT**

**Hardik Makkar-** conceptualization, data curation, investigation, formal analysis, methodology, visualization, writing - original draft & editing.

**Nghi Tran-** Investigation, methodology, formal analysis

**Yu-Chang Chen-** Investigation, methodology, formal analysis

**Kang I Ko-** conceptualization, supervision, resources, Investigation

**Rebecca G. Wells-** conceptualization, supervision, visualization, writing

**Kyle H. Vining** conceptualization, methodology, funding acquisition, supervision, visualization, writing, resources

## Data Availability Statement

The data that support the findings of this study will be openly available in Dyrad with a DOI after publication.

