## Supporting Information for "Matrix Stiffness Governs Fibroblast-Driven Immune Homeostasis in Gingival Tissues"

### Methods

#### Isolation of Donor-Derived Gingival Fibroblasts

Gingival fibroblasts were isolated from healthy human gingival tissue from the University of Pennsylvania, Periodontology Clinic (IRB exemption #844933, PI: Ko). Freshly obtained gingival tissues were washed in 1x HBSS without calcium and magnesium supplemented with antibiotic-antimycotic (1X, Gibco). The gingival tissues were cut into small pieces and placed in a T25 flask for 72 hours undisturbed and supplemented with Dulbecco's modified eagles medium (low glucose;1g/L) with 10% FBS and antibiotic-antimycotic (1X, Gibco). Cells were expanded, cryopreserved, and passages 2-4 were used for experiments.

#### G-CSF Mobilized CD34+ HSPC Cell Culture and Differentiation

Cells obtained from Fred Hutch Cancer Center were cultured at 20,000 cells/well 5 days per passage in serum-free StemSpan medium II (Stem Cell Technologies) supplemented with 100 ng/mL of stem cell factor (SCF; Peprotech), 100 ng/mL of Flt-3 ligand (FL3TL; Peprotech), 50 ng/mL of thrombopoietin (TPO; Peprotech), 20 ng/mL of IL-3 (Peprotech), and 700 nM UM729 (Stem Cell 15 Technologies). Cells were seeded into 96-well U-bottom plates with 200 µL of prewarmed media. The cultures were maintained at 37°C in a humidified atmosphere of 5% CO2. Half media changes were performed with freshly supplemented cytokines every 3 days. Expanded HSPCs at passage 2 (10 days total) were used for differentiation. Cells were differentiated for 7-days at a density of $1 \times 10^5$ cells/mL in Iscove's modified Dulbecco's medium (IMDM) containing 10% heat-inactivated FBS, 25 mmol/ L Hepes, 2 mmol/L L-glutamine, 0.1 mmol/L nonessential amino acids, 100 U/mL penicillin, and 100 µg/mL streptomycin (GIBCO), supplemented with 100 ng/mL of granulocyte-macrophage colony-stimulating factor (GM-CSF; Peprotech), 100 ng/mL of Flt-3 ligand (FL3TL; Peprotech) in 96 well U bottom plates (Costar, Cambridge, MA) at 37°C in a humidified atmosphere of 5% CO2. Media changes were performed every 3 days with freshly supplemented cytokines. Cells were cryopreserved in BamBanker (Fujifilm) at 7 days of myeloid differentiation.

#### Preparation of Gingival ECM hydrogels

Gingival ECM hydrogels were prepared as described previously[1-4]. Briefly, alginate (Pronova) was dissolved in a solution of 1x HBSS without calcium nor magnesium (14175095, Gibco) with HBSS/HEPES buffer) to a final w/v concentration of 5%. On the day of IPN fabrication, in a glass vial with a magnetic stir bar, bovine telo collagen I (Advanced Biomatrix) was neutralized on ice in v/v ratio of 17:3 with a solution created from 10x HEPES (14180046, ThermoFisher), 1 M HEPES, 1x HBSS, and 1M sodium hydroxide (NaOH) mixed in a v/v ratio of 10:2:2:1, respectively. Extra NaOH was added dropwise until the phenol red indicated a pH of ~7. HBSS/HEPES buffer was used to adjust the final concentration of collagen to 4 mg/mL. Subsequently, a 100 mg/mL CaCO3 slurry was prepared in endotoxin-free ultra-pure water. With these solutions ready, an aliquot of the CaCO3 slurry was incorporated into the neutralized collagen at a final concentration of 10, 20, or 30 mM to achieve storage moduli of 800 to 2000 Pa, respectively. Then, alginate was added to the mixture at a final w/v concentration of 1%. These solutions were mixed with a magnetic stirrer. Next, a solution of 0.4 g/mL glucono-delta-lactone was formulated with HBSS/HEPES buffer immediately before its incorporation into the gel mixture at a 4x molar excess of CaCO3. The solution was vigorously stirred and mixed by pipetting. Working quickly, the liquid hydrogel solution was cast in 96 (60 $\mu$L) or 48 well-plates (150 $\mu$L) and incubated at 37°C for further experimentation.

#### Rheological Characterization of hydrogels and gingival tissues

The rheological properties of the gingival ECM hydrogels were measured on the AR-G2 rheometer (TA Instrument) as shown previously[4]. 100 $\mu$L of the hydrogel solution was aliquoted onto the rheometer loaded with a 20 mm flat plate at 37 °C. Excess solution was wiped off, and a solvent trap (sealed with mineral oil) was added to prevent evaporation during the rheometric tests. At an oscillation of 1 Hz and strain of 1%, the storage modulus (G') and loss tangent (tan δ) were recorded until equilibrium was achieved (1 to 2 h), after which a frequency sweep was performed (0.1 to 10 Hz, 1% strain). For the rheological measurements of human gingival tissues, samples are placed on the Peltier-controlled chamber at 37°C submerged in HBSS/HEPES solution with 0.01N compressive preload. Viscoelastic properties such as storage modulus and tan(delta) are measured using a frequency sweep from 0.01-25 Hz with 0.5% strain, followed by a stress relaxation test with 10% strain.

**Cell culture and hydrogel encapsulation**

To prevent collagen contraction, surface modification of 96 and 48 flat-bottom culture plates was done with a 2 mg/ml dopamine solution (Sigma Aldrich, USA) prepared in 10 mM Tris-HCl (pH 8.5) for 2h at room temperature[5]. After 2 hours of incubation, the dopamine solution was gently aspirated, and the surfaces were washed twice with sterile filtered deionized (DI) water to remove unbound dopamine. The washed substrates were kept sterile at room temperature until use. Gingival Fibroblasts (GFs) were isolated from healthy donor gingival tissues, cultured as explants. The isolated cells were cultured in Dulbecco's modified Eagle's medium (low glucose;1g/L) with 10% FBS. GFs suspended in HBSS/HEPES buffer and added to the collagen mixture, prior to the addition of alginate, at a final concentration of 1E06 cells/mL. Once all the hydrogel components were added and mixed, 60 $\mu$L or 150 $\mu$L hydrogel constructs were prepared in a 96- or 48-well flat-bottom well plates. The plates were then moved to a cell incubator at 37°C and 5% CO2. After 15 and 30 minutes, HBSS/HEPES buffer was added to the hydrogels, respectively, to maintain a pH ~7. At 30 minutes, the buffer in the wells containing the ECM hydrogels was replaced with a low-serum defined media[6].

**Exposure of gingival ECM hydrogels to pattern recognition receptor agonists**

Gingival ECM hydrogels cultures for four days were exposed with TLR-2 agonist (Pam3CSK4, Invivogen), TLR-4 agonist (ultrapure *E. coli* LPS, Invivogen), TLR-3 agonist and TLR-9 agonist for 24 h. Following challenge, the culture supernatants were collected and stored at –80°C for downstream cytokine analysis. The concentrations are summarized in **Supplementary Table T2.**

**Inhibition studies on Gingival ECM hydrogels**

For pharmacological inhibition studies, the inhibitors were added to the culture media following encapsulation. The concentrations used for each inhibitor are summarized in **Supplementary Table T2.**

**Retrieval of cells from ECM hydrogels and Viability assessment**

Encapsulated cells were recovered by enzymatic digestion of the hydrogels[1-3,7]. Briefly, hydrogels were incubated with a digestion solution containing 300 U/mL collagenase type I (Thermo Scientific) and 34 U/mL alginate lyase (Sigma-Aldrich) in Dulbecco's phosphate-buffered saline (DPBS) supplemented with calcium and magnesium, and 0.5% bovine serum albumin (BSA), at 37°C for 15 minutes. An additional 100–150 $\mu$L of 300 U/mL collagenase I was then added, and the gel was pipetted up and down using a P1000 pipette, followed by a second incubation at 37°C for 40 minutes. To terminate digestion, 0.5 mL of wash buffer (DPBS without calcium and magnesium, supplemented with 2 mM EDTA and 0.5% BSA) was added. Cells stuck to the bottom of the plate were detached using ice-cold PBS -/- and EasySep buffer washes (Stem Cell Technologies). Additional washes were performed if cells remain attached, or if undigested hydrogel was visible. The cell suspension was transferred to a v-bottom 96-well plate and centrifuged at 400 × g for 5 minutes at 4°C. The supernatant was removed, and the pellet was washed once more with 200 $\mu$L of cold wash buffer on ice to obtain a clean cell pellet. Cell viability and total cell number were assessed using Trypan blue Assay using Countess™ automated cell counters.

**Flow cytometry**

Cells recovered from hydrogels were first stained with a fixable viability dye (Thermo Fisher Scientific) to exclude dead cells. Following viability staining, Fc receptors were blocked using human TruStain Fc receptor blocking reagent, and cells were stained with a panel of anti-human monoclonal antibodies targeting surface markers relevant to mesenchymal stromal cells, hematopoietic stem cell (HSC) expansion, myeloid differentiation, and immune activation (**Supplementary table T3)**. All antibodies were obtained from BioLegend. Staining was performed in eBioscience™ Flow Cytometry Staining Buffer (Thermo Fisher Scientific) according to the manufacturer's instructions. After staining, cells were fixed with 2% paraformaldehyde for 20 minutes at room temperature, followed by three washes with FACs buffer. Fixed samples were stored at 4°C in the dark and analyzed within 48 hours. Single-color controls were prepared using compensation beads (BioLegend) for spectral unmixing. Data acquisition was performed using a 5-L Cytek® Aurora spectral flow cytometer at the Children's Hospital of Philadelphia (CHOP) Flow Cytometry Core. Data were analyzed using FCS Express.

**Enzyme Linked Immunosorbent Assay (ELISA) for secretome analysis**

In accordance with manufacturer's instructions, ELISAs for cytokines IL-6, IL-8, CCL-2 (all from Biolegend) were performed using the culture supernatants. The absolute cytokine values were normalized to the total protein content using BCA assay (Thermo Fisher Scientific).

**Immunostaining of cryosections and whole mounts immunostaining.**

The Gingival ECM hydrogels were fixed in 4% PFA (EMS), processed, and embedded in optimum cutting temperature compound (Tissue-Tek). Tissue sections (10 $\mu$m thick). For immunostaining[8], the sections were first washed with DPBS, antigen retrieval with 0.5% Triton-X followed by blocking of nonspecific staining. The sections were incubated with respective primary antibodies overnight at 4°C, washed and incubated with respective secondary antibodies for 2 hours at 4°C (**Supplementary table T3)**. The nuclei were counterstained with DAPI and mounted with anti-fade fluorescent mounting medium (Abcam). The slides were visualized under confocal microscope (Leica Stellaris, Leica Microsystems equipped with Leica Application Suite X software). Whole mount immunostaining was done as previously described[9]. Briefly, antigen retrieval and permeabilization was done by immersing the hydrogels in 0.5% Triton-X in phosphate-buffered saline (PBS) under orbital shaking for 2 h. The gels were then incubated in blocking solution (PBS containing 0.5% Triton-X, 10% goat serum, and 10% bovine serum albumin) overnight under orbital shaking. Tissues were incubated with primary antibodies (**Supplementary table T3)** for 2 d at 4 °C followed by washing under orbital shaking for 3 h. This was followed by incubation with respective secondary antibodies (**Supplementary table T3)** for 2 d at 4 °C and counterstaining with DAPI. Whole mounts were visualized using laser scanning confocal microscopy (Leica Stellaris, Leica Microsystems). Image analysis was done using Fiji (NIH) and Imaris software (Oxford Instruments).

**Second Harmonic Generation Imaging and image analysis.**

Second harmonic generation (SHG) imaging of formalin fixed paraffin embedded sections of human gingiva and viable whole gingival tissues were collected on Leica SP8-MP Upright with an external non-descanned hybrid detector (HyD – RLD2). The laser was tuned to 910nm, and the filter associated with the lens was 435~485nm. All images were processed from forward and backward SHG signals using z-stacks in FIJI (ImageJ) and Imaris software (Oxford Instruments). Briefly, to quantify the orientation of ECM collagen fibers, SHG images were analyzed using directionality plugin in FIJI/ImageJ[10,11]. This plugin computes a histogram that represents the data as a percentage of pixels in a given direction (between 1 to 180 degrees). The resultant direction of orientation and the pixel percentage counts are imported into Prism (Dotmatics). The pixel percentage counts for orientation of collagen fibers in a single layer are presented as a frequency histogram and that for the whole *z*-stack are presented as a heat map. Further, The CT-FIRE program (http://loci.wisc.edu/software/ctfire, version 1.3, beta 267) was utilized to quantify parameters of individual collagen fibers width, and straightness, in each SHG image.

**Cell Viability analysis**

The viability of the GFs in hydrogels was determined using Lactate Dehydrogenases (LDH) Assay (Promega) In accordance with manufacturer's instructions.

**Phagocytosis Assay**

Phagocytic activity was assessed using a fluorescent bioparticle uptake assay. Hydrogel retrieved cells were seeded at a density of $1 \times 10^4$ cells per well into glass bottom plates. Plates were incubated at 37 °C with 5% $CO_2$ for at least 1 hour to allow cells to settle. Fluorescently labeled bioparticles (e.g., pH-sensitive *E. coli* conjugates) were thawed, diluted 20X and 100 $\mu$L of the prepared bioparticle suspension was added. Cells were then incubated for 90 minutes at 37 °C to allow for phagocytosis and intracellular acidification. The plates were imaged using confocal microscope (Leica Stellaris, Leica Microsystems) and image analysis was done using Fiji (NIH).

**Bulk RNA- sequencing and analysis**

Cells encapsulated in hydrogels were harvested and lysed using RLT buffer supplemented with 1% β-mercaptoethanol (β-ME) (Sigma-Aldrich), following standard protocols for disruption and homogenization of cell pellets. Briefly, cell pellets were resuspended in 350 $\mu$L of RLT + β-ME and pipetted vigorously to ensure complete lysis and homogenization. Lysates were stored at –80°C or processed immediately for RNA extraction. Total RNA was isolated using a commercial column-based purification kit (RNeasy Mini Kit, Qiagen), according to the manufacturer's instructions. This included an on-column DNase I digestion step to remove genomic DNA contamination and elution in RNase-free water. RNA concentration and integrity were assessed by spectrophotometry (NanoDrop) prior to sequencing. Bulk RNA sequencing was performed by Azenta Life Sciences. Library preparation, quality control, sequencing, and initial data processing were conducted by Azenta. Differential gene expression results from bulk RNA sequencing were analyzed using Reactome pathway enrichment to identify significantly altered biological pathways. Enrichment analysis was performed using the Reactome Pathway Database (https://reactome.org) by submitting the list of significantly upregulated and downregulated genes. Overrepresentation analysis was carried out using the Reactome analysis tool, which applies a hypergeometric test to assess pathway enrichment, followed by false discovery rate (FDR) correction for multiple testing. Pathways with an FDR < 0.05 were considered significantly enriched. The top enriched up- and downregulated pathways were visualized based on normalized enrichment scores and *p*-values to highlight key biological processes altered in the dataset. For a dedicated analysis of gene sets, z-score normalized gene counts were plotted as heatmaps. In parallel, Gene Ontology (GO) enrichment analysis was performed using the NIH DAVID Bioinformatics Resources (v6.8). Official gene symbols were uploaded, and functional annotation clustering was conducted. Enrichment significance was assessed using DAVID's modified Fisher Exact test (EASE score), with multiple testing correction by the Benjamini–Hochberg method. GO biological process terms with adjusted $p < 0.05$ were considered significantly enriched.

**Human gingival explants culture and transglutaminase injection**

Deidentified human gingival tissues were obtained from the University of Pennsylvania Periodontology Clinic (IRB #844933, PI: Ko). Immediately following collection, explants were washed with HBSS buffer (without calcium and magnesium) supplemented with 1% antibiotic–antimycotic solution. Transglutaminase (10 U; SAE0159, Sigma) reconstituted in PBS was injected into the tissues using a hypodermic syringe; control samples received PBS alone. Tissue punches (2 mm) were prepared using a sterile biopsy punch and placed in transwell inserts under air–liquid interface conditions. Explants were cultured for 3 days in low-serum medium containing ascorbic acid (10 mg ml–1), hydrocortisone (50 $\mu$g ml–1), basic fibroblast growth factor (10 ng ml–1), vascular endothelial growth factor (20 ng ml–1), epidermal growth factor (5 ng ml–1), 1% penicillin–streptomycin, and SITE supplement (selenium, 5 ng ml–1; insulin, 10 $\mu$g ml–1; transferrin, 5.5 $\mu$g ml–1; ethanolamine, 2 $\mu$g ml–1). During the final 24 h of culture, explants were treated with a TLR2 agonist (Pam3CSK4, 1 $\mu$g ml–1). Following culture, tissues were subjected to rheological measurements, and conditioned media were collected for ELISA analysis.

**Supplementary Figures**

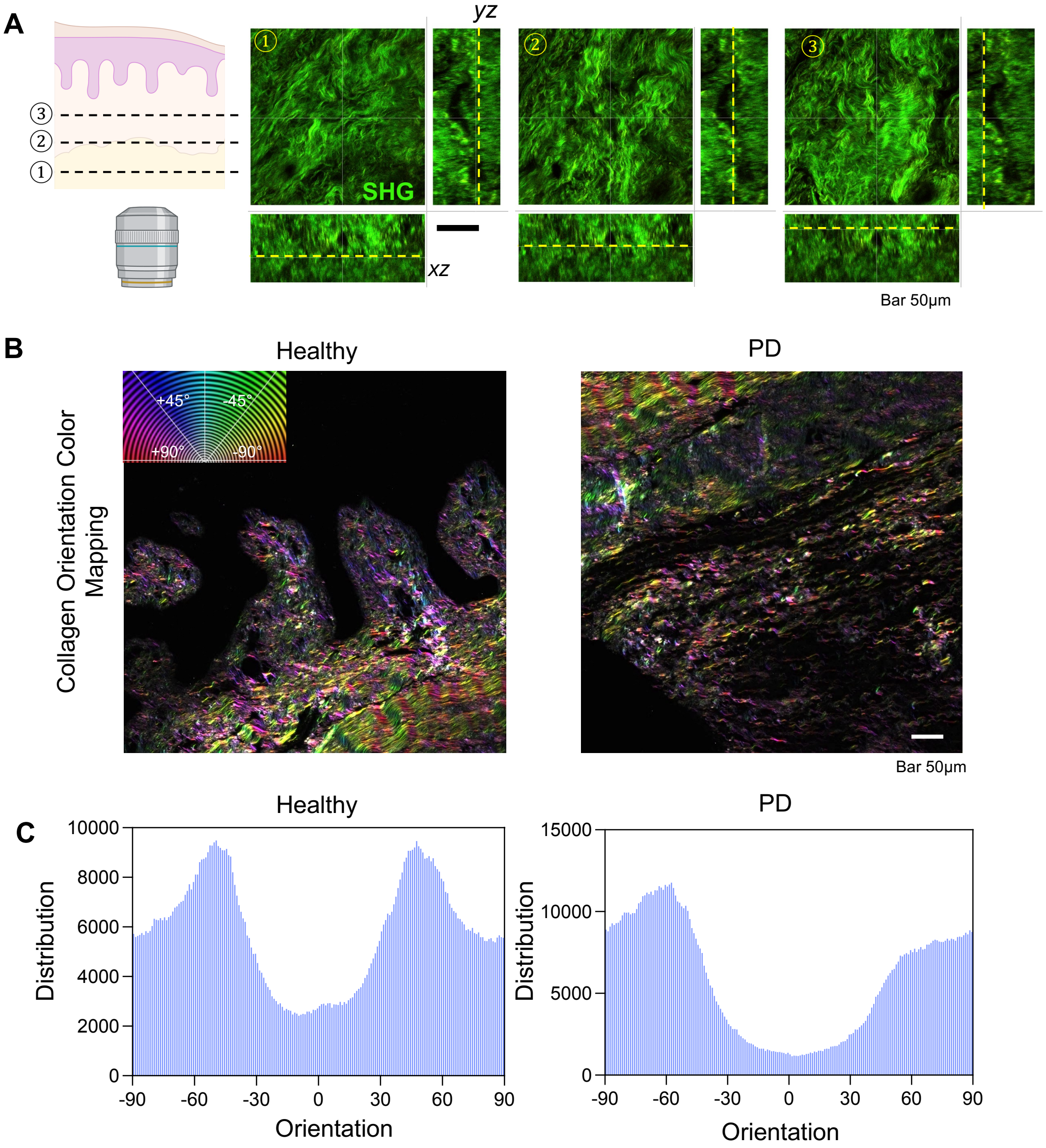


**Supplementary Figure S1. (A)** z-sections of whole-mount SHG image of gingival connective tissue. **(B)** Collagen orientation color map of collagen fibers in healthy and diseased gingival connective tissue. **(C)** The angular distribution histogram for the healthy and diseased site (PD). Scale bars are as indicated.

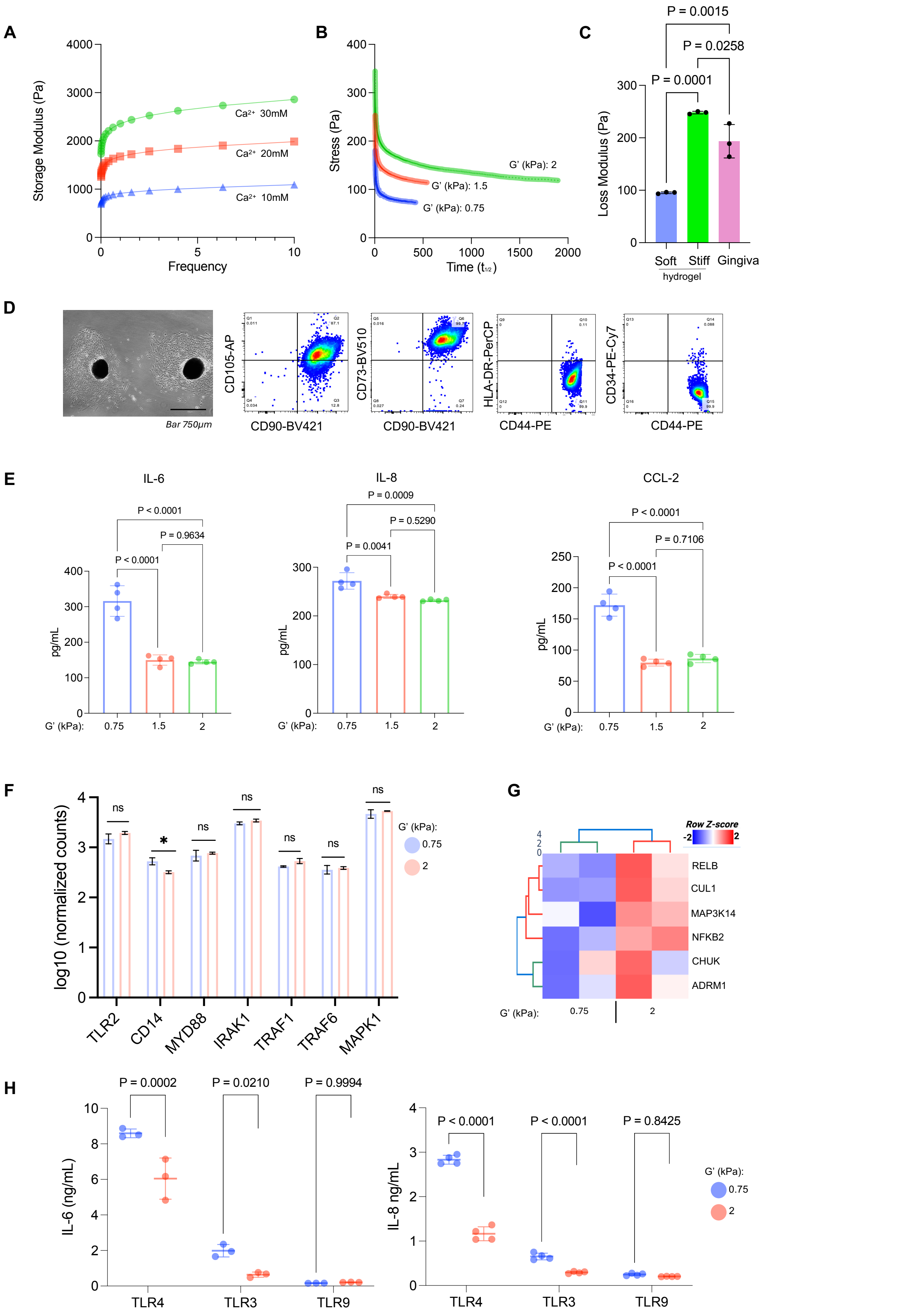


**Supplementary Figure S2.** **(A)** Oscillatory shear rheology- frequency sweep and **(B)** stress relaxation behavior of gingival ECM hydrogels **(C)** Loss modulus of gingival hydrogels and healthy gingival tissue. **(D)** Isolation of GFs from healthy donors and culture as explants. Flow cytometry confirms the mesenchymal stromal surface marker profile, showing cells are positive for CD105, CD73, and CD90, and negative for hematopoietic and endothelial markers HLA-DR, CD44, and CD34. **(E)** ELISA quantification of secreted IL-6, IL-8, and CCL-2 from GFs cultured in hydrogels of varying stiffness. **(F)** Gene expression analysis of key components of the TLR2 signaling pathway in gingival fibroblasts (GFs) cultured in soft versus stiff hydrogels. **(G)** Heatmap of differentially expressed genes in non-canonical NF-κB pathway in GFs cultured in soft and stiff hydrogels. **(H)** Secretion of IL-6 and IL-8 by GFs encapsulated in soft and stiff hydrogels exposed with TLR3, TLR4 & TLR9 agonists. Scale bars are as indicated. Data are shown as mean ± SD. P-values from statistical tests are indicated.

**A**

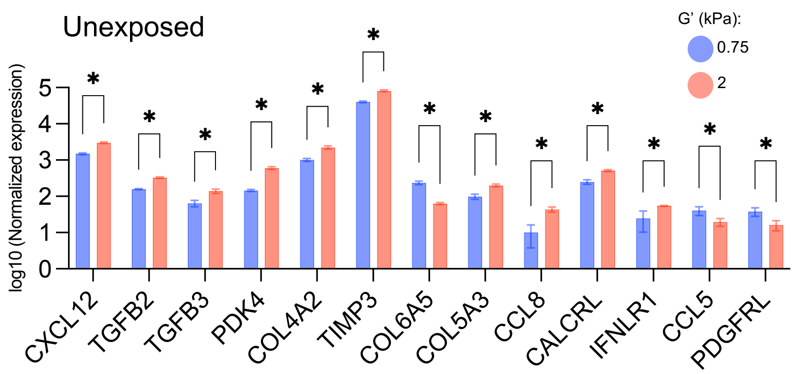


| Gene | P value | q value |
|---|---|---|
| CXCL12 | 0.000101 | 0.000286 |
| TGFB2 | 0.000025 | 0.000170 |
| TGFB3 | 0.006385 | 0.009684 |
| PDK4 | 0.000025 | 0.000170 |
| COL4A2 | 0.000521 | 0.001185 |
| TIMP3 | 0.000105 | 0.000286 |
| COL6A5 | 0.000067 | 0.000286 |
| COL5A3 | 0.002407 | 0.004107 |
| CCL8 | 0.029657 | 0.033734 |
| CALCRL | 0.001482 | 0.002890 |
| IFNLR1 | 0.047410 | 0.049780 |
| CCL5 | 0.025458 | 0.031592 |
| PDGFRL | 0.019543 | 0.026677 |

**B**

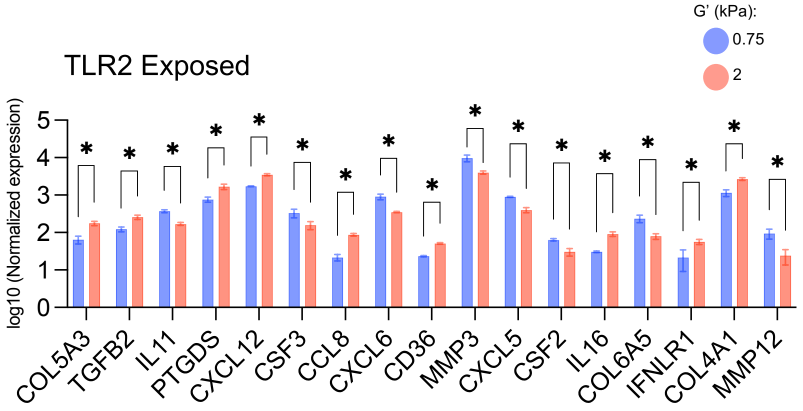


| Gene | P value | q value |
|---|---|---|
| COL5A3 | 0.002093 | 0.004584 |
| TGFB2 | 0.003592 | 0.005829 |
| IL11 | 0.000690 | 0.002462 |
| PTGDS | 0.006381 | 0.009492 |
| CXCL12 | 0.000244 | 0.001088 |
| CSF3 | 0.031890 | 0.033484 |
| CCL8 | 0.000175 | 0.001088 |
| CXCL6 | 0.003432 | 0.005829 |
| CD36 | 0.000062 | 0.001088 |
| MMP3 | 0.007897 | 0.010069 |
| CXCL5 | 0.000234 | 0.001088 |
| CSF2 | 0.002311 | 0.004584 |
| IL16 | 0.001201 | 0.003063 |

**Supplementary Figure S3**. Normalized expression of genes in gingival fibroblasts encapsulated in soft and stiff ECM hydrogels without **(A)** and with **(B)** TLR2 agonist. All data are shown as mean ± SD. P-values and q values from statistical tests are indicated in tables.

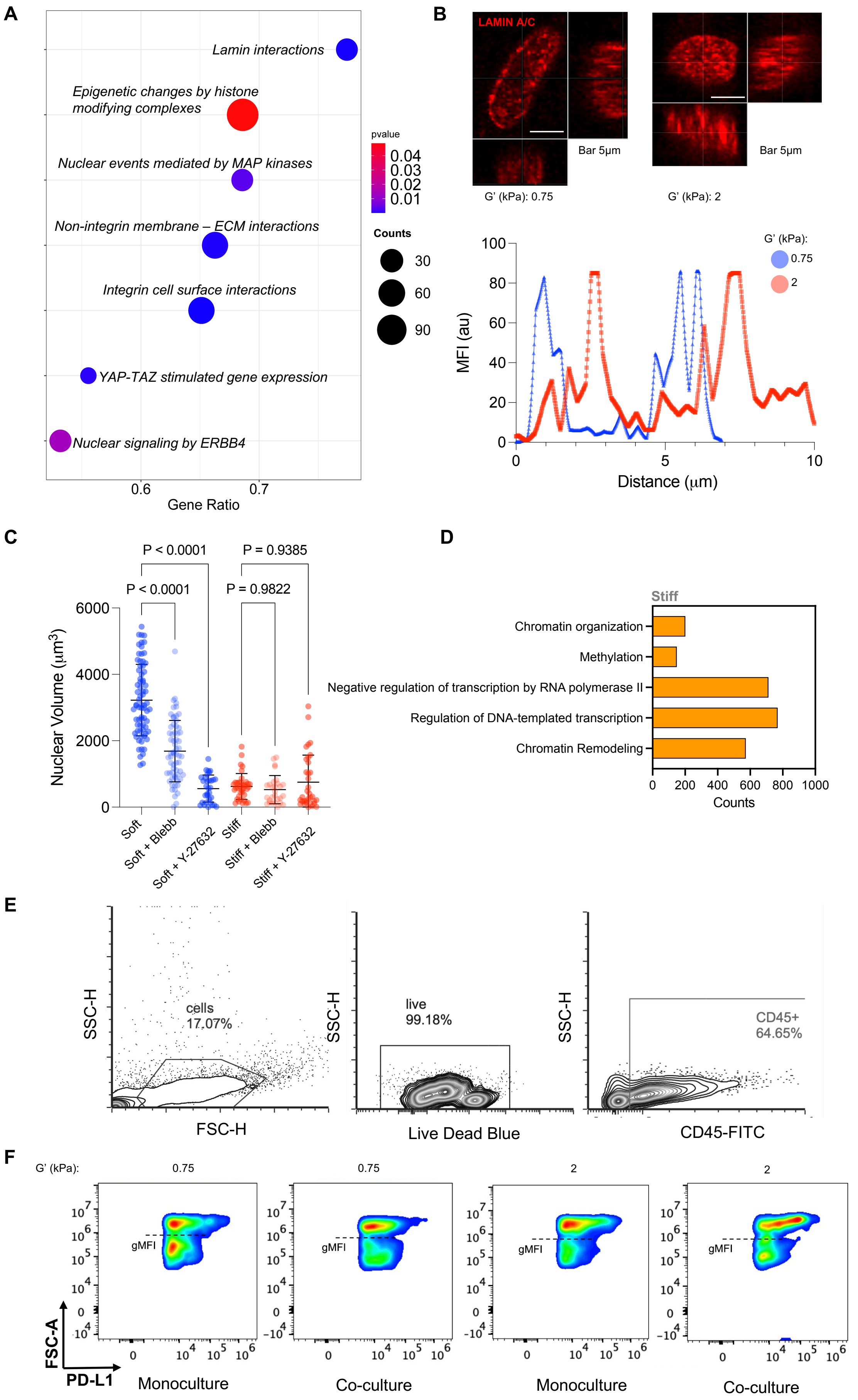


**Supplementary Figure S4**. **(A)** A bubble plot illustrates the enrichment of pathways associated with nuclear and epigenetic changes in GFs cultured in stiff (2kPa) hydrogels. **(B)** Immunofluorescence images show the distribution of Lamin A/C (red)

and the cell nucleus (stained with DAPI, blue) in cells on soft and stiff substrates. The line scan profile below the images quantifies the fluorescence intensity of Lamin A/C across the nucleus, showing a more peripheral localization of lamin AC in GFs encapsulated in stiff hydrogels compared to more diffuse expression under soft condition. **(C)** Plots representing the nuclear volume of GFs in soft and stiff hydrogels treated with myosin II inhibitor Blebbistatin or the ROCK inhibitor Y-27632. **(D)** Plot showing the enrichment of Gene Ontology (GO) terms related to epigenetic regulation of GFs in stiff hydrogels. **(E)** Gating strategy for myeloid cells and gingival fibroblasts co-cultured in gingival ECM hydrogels. **(F)** Representative flow cytometry plots showing the expression of PD-L1 on myeloid cells (gated on live, CD45+ cells) after culture in soft or stiff ECM hydrogels, either in monoculture or in co-culture with gingival FBs. Scale bars are as indicated. Data are shown as mean ± SD. P-values from statistical tests are indicated.

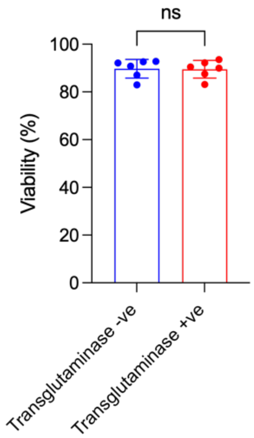


**Supplementary Figure S5**. Viability of gingival explants ex vivo with and without transglutaminase treatment. All data are shown as mean ± SD. P-values from statistical tests are indicated

**Supplementary Table T1.** Formulations of artificial ECM hydrogels.

| Hydrogel | Collagen (mg/mL) | VLVG alginate (% w/v) | $CaCO_3$ (% w/v) | GDL (mM) |
|---|---|---|---|---|
| Soft viscous | 4 | 1 | 0.10 | 40 |
| Stiff viscous | 4 | 1 | 0.30 | 120 |

**Supplementary Table T2.** Pharmacological inhibitors and cytokine treatments.

| Name | Synonyms | Target | Concentration | Manufacturer |
|---|---|---|---|---|
| Pam3CSK4 | TLR2 ligand | TLR2 | 1 µM | Invivogen |
| Poly(I:C)HMW | TLR3 ligand | TLR3 | 10 ug/mL | Invivogen |
| ODN-M362 | TLR9 ligand | TLR9 | 5 µM | Invivogen |
| IKK inhibitor | BMS-345541 | IKK | 10 µM | Selleckchem |
| Decitabine | DNMT inhibitor | DNMT | 0.5, 1 µM | Selleckchem |
| Blebbistatin | | NMII | 25 µM | Selleckchem |

| | | | | |
|---|---|---|---|---|
| ROCK inhibitor | Y-27632 | ROCK | 25 µM | Selleckchem |
| LPS | LPS | TLR4 agonist | 1 ug/mL | Invivogen |
| IFN-gamma | IFNgamma | activates TNFalpha | 100 ng/mL | Peprotech |

**Supplementary Table T3.** Details of primary and secondary antibodies used in the study

| Antibody | Specification | Dilution | Source |
|---|---|---|---|
| Phalloidin i-Fluor | | 1:1000 | Abcam (ab176753) |
| Lamin A/C (4C11) | Mouse monoclonal | 1:400 | Cell Signaling Technologies (4777T) |
| Phospho RelB (Ser552) | Rabbit monoclonal | 1:200 | Cell Signaling Technologies (4999S) |
| DNMT1 (D63A6) | Rabbit monoclonal | 1:200 | Cell Signaling Technologies(5032T) |
| Secondary antibody | Alexa Fluor-488 (Goat anti-mouse) | 1:250 | Molecular Probes (#A11001) |
| Secondary antibody | Alexa Fluor-594 (Goat anti-rabbit) | 1:300 | Molecular Probes (#A11037) |
| Brilliant Violet 421™ anti-human CD90 | Mouse monoclonal | 1:100 | Biolegend (328121) |
| APC anti-human CD105 | Mouse monoclonal | 1:20 | Biolegend (323207) |
| PerCP anti-human HLA-DR | Mouse monoclonal | 1:100 | Biolegend (307627) |
| PE/Cyanine 7 anti-human CD34 | Mouse monoclonal | 1:100 | Biolegend (343515) |
| PE anti-human CD44 | Mouse monoclonal | 1:100 | Biolegend (397503) |
| Brilliant Violet 421™ anti-human CD73 | Mouse monoclonal | 1:100 | Biolegend (344043) |
| PE/Dazzle™ 594 anti-human PD-L1 | Mouse monoclonal | 1:100 | Biolegend (329731) |
| Brilliant Violet 711™ anti-human CD11b | Mouse monoclonal | 1:100 | Biolegend (301343) |

**Supplementary Table T4 – Statistical tests in data of main figure**

| Data | Test | Passed Normality test? |
|---|---|---|
| **Fig. 1**I | Ordinary one-way ANOVA<br>Dunnett's multiple comparisons test | Yes |
| **Fig.** 2D | Ordinary one-way ANOVA<br>Tukey’s multiple comparisons | Yes |
| **Fig.** 3C | Ordinary one-way ANOVA<br>Tukey’s multiple comparisons | Yes |
| **Fig.** 4B | Unpaired t test, two-tailed | Yes |
| **Fig.** 4D | Two-way ANOVA<br>Šídák's multiple comparisons test | Yes |
| **Fig.** 4E | Ordinary one-way ANOVA<br>Tukey’s multiple comparisons | Yes |
| **Fig.** 4H | Unpaired t test, two-tailed | Yes |
| **Fig.** 4I | Two-way ANOVA<br>Šídák's multiple comparisons test | Yes |
| **Fig.** 5C | Two-way ANOVA<br>Šídák's multiple comparisons test | Yes |
| **Fig.** 5D | Unpaired t test with Welch's correction | Yes |
| **Fig.** 5E | Ordinary one-way ANOVA<br>Tukey’s multiple comparisons | Yes |
| **Fig.** 6B | Two-way ANOVA<br>Uncorrected Fisher's LSD | Yes |
| **Fig.** 6C | Unpaired t test, two-tailed | Yes |
| **Fig.** 6E | Paired t test, two-tailed | Yes |
| **Fig.** 6F | Paired t test, two-tailed | Yes |

**AUTHENTICATION OF KEY BIOLOGICAL AND/OR CHEMICAL RESOURCES**

1) All acquired compounds and reagents were authenticated for both identity and purity, based on certificate of analysis from manufacturers.

2) The fluorescent probes, laser and filter parameters related to the confocal microscopy have been extensively tested and selected to minimize potential cross talk between different excitation and emission channels
3) Primary human cells for in vitro experimentation were routinely tested for mycoplasma and bacterial contaminations.
